# Mitochondrial COX4I2 drives pericyte-dependent inflammation and emphysema

**DOI:** 10.64898/2026.02.09.703513

**Authors:** Claudia F. Garcia Castro, Vidya Srokshna Balasubramanian Lakshmi, Stefan Hadzic, Claudio Nardiello, Rolf D. Glaser, Maik Hüttemann, Lawrence I. Grossman, Baktybek Kojonazarov, Muchen Li, Sandipan Jash, Janine Koepke, Marija Gredic, Cheng-Yu Wu, Luca Giordano, Matthias Hecker, Christos Samakovlis, Edma Loku, Anis Cilic, Julian Better, Ulrich Matt, Brigitte Mueller, Knut Stieger, Lyubomyr Lytvynchuk, Learta Pervizaj-Oruqaj, Andreas Guenther, Jochen Wilhelm, Susanne Herold, Slaven Crnkovic, Grazyna Kwapiszewska, Michael P. Murphy, Friedrich Grimminger, Marek Bartkuhn, Werner Seeger, Norbert Weissmann, Oleg Pak, Natascha Sommer

**Author notes:** shared first authorship (Claudia F. Garcia Castro, Vidya Srokshna Balasubramanian Lakshmi). shared last authorship (Oleg Pak, Natascha Sommer).

## Abstract

Chronic obstructive pulmonary disease (COPD) is characterized by neutrophilic inflammation, emphysema, and mild pulmonary hypertension (PH). Oxidative/nitrosative stress are key drivers, but specific mitochondrial mechanisms remain unclear. We show increased expression of the regulatory mitochondrial cytochrome c oxidase subunit 4 isoform 2 (COX4I2) in an early murine model and human COPD. After 8 months of cigarette smoke exposure, *Cox4i2^−/−^* mice were completely protected from emphysema but not from PH, associated with reduced nitrosative stress, inflammation, and apoptosis. Using a novel *Cox4i2* reporter mouse and in situ hybridization of human lungs, COX4I2 was detected in precapillary ACTA2^+^ cells and capillary pericytes. COX4I2 promotes mitochondrial reactive oxygen species (mtROS) production in these cells, thereby enhancing neutrophil migration and alveolar type II cell apoptosis, and modulates angiogenesis. In contrast to *Cox4i2^−/−^*, mitochondria-targeted antioxidant MitoQ reversed emphysema and PH, suggesting pericyte-specific regulation of COPD pathologies and mtROS inhibition as a therapeutic approach in COPD.

**Graphical abstract:** 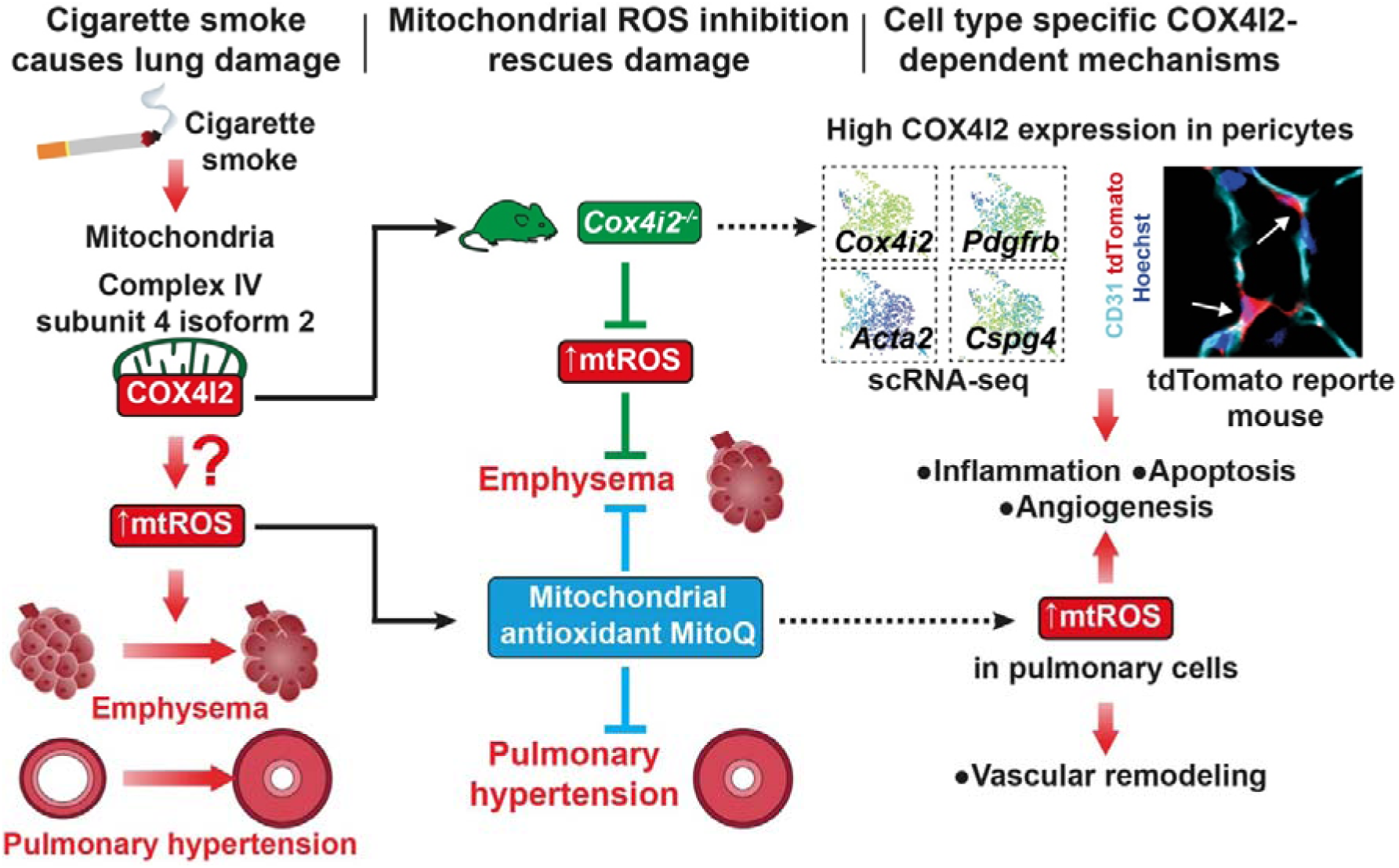

## Introduction

Chronic obstructive pulmonary disease (COPD) ranks as the third leading cause of death worldwide^1^ and is predominantly caused by tobacco-associated or environmental smoke particles and gases in combination with developmental and genetic risk factors^2–5^. COPD is a heterogeneous lung disease characterized by structural alterations of the airways (bronchitis and bronchiolitis), alveoli (emphysema) and pulmonary vasculature (pulmonary hypertension, PH)^2,5,6^. Underlying pathomechanisms include predominantly neutrophilic inflammation, alveolar apoptosis and abnormal proliferation of smooth muscle cells (SMCs) and fibroblasts^6^. However, the cell type specific impact of these various cells in the underlying compartmentalised pathology is not completely elucidated. Oxidative stress originating from internal and external sources in COPD promotes chronic inflammation, cellular senescence and apoptosis and impairs DNA repair mechanisms^7^.

Previously, we demonstrated that a murine model of chronic cigarette smoke (CS) exposure replicates the development of PH after 3 months of CS exposure and the development of emphysema after 8 months, with both alterations persisting even after cessation of CS exposure^8,9^. In this model, we demonstrated that superoxide derived from the NADPH oxidase-containing subunit NOXO1, along with the excessive release of nitric oxide (NO) triggered by the increased expression of inducible nitric oxide synthase (iNOS), result in nitrosative stress. Nitrosative stress was indicated by the generation of 3-nitrotyrosine, a biomarker for the formation of the oxidant peroxynitrite from NO and superoxide. These findings were consistent with the suggested role of nitrosative stress in the development of CS-induced emphysema and PH^8,9^. Importantly, iNOS expression in macrophages specifically regulates CS-induced PH but not emphysema, emphasizing the distinct contributions of different cell types to different CS-induced pathological changes caused by different sources of ROS and NO that participate in oxidative and nitrosative stress^10^.

Mitochondria may be sources of ROS in addition to NOXO1-dependent NADPH oxidases in different pulmonary cell types^11^. CS has been shown to increase mitochondrial ROS (mtROS) production in human primary alveolar type II cells (ATIIs) and human bronchial epithelial cells (NHBE)^12^. However, the *in vivo* relevance, cellular source and mechanism of mtROS production for the development of CS-induced inflammation, emphysema and PH are unknown. The main components of CS, such as cyanide and carbon monoxide, may inhibit mitochondrial cytochrome *c* oxidase (COX) activity and increase mitochondrial oxidative damage and mtROS generation^13^, as shown previously by our laboratory for acute hypoxia^14,15^. Specifically, we demonstrated that the lung-specific isoform 2 of the cytochrome *c* oxidase subunit 4 (COX4I2) of mitochondrial complex IV, which catalyses the final step of the electron transport chain (ETC), is essential for increased mitochondrial superoxide release from complex III in precapillary pulmonary arterial SMCs (PASMCs) when COX activity decreases even during mild hypoxia^14,15^. Moreover, mtROS production from complex III may be involved in CS-associated immune cell activation via toll like receptor 4 (TLR4) signalling^16,17^. In this context, it has been suggested that increased COX4I2 expression contributes to a CS-induced increase in the contribution of COX activity to inflammation, although the exact mechanism by which alterations in COX affect mitochondrial function or inflammation has not been investigated^18^. We thus hypothesized that COX4I2-dependent mtROS generation and/or oxidative stress may contribute to CS-induced nitrosative stress, inflammation, emphysema and PH and that inhibition of mtROS generation and/or oxidative stress may serve as a novel therapeutic approach.

## Results

### COX4I2 is upregulated in early COPD and contributes to CS-induced nitrosative stress

Previously, we showed that iNOS-mediated NO release^8^ and superoxide originating from NOXO1-containing NADPH oxidases^9^ contribute to generation of the highly reactive oxidant peroxynitrite and thereby to CS-induced nitrosative stress, indicated by increased levels of 3-nitrotyrosine formation, which was associated with the development of CS-induced emphysema and PH. We now further hypothesize that mtROS release facilitated by the expression of the COX subunit COX4I2 contributes to CS-induced nitrosative stress by leading to peroxynitrite formation, 3-nitrotyrosine production, oxidative stress and pulmonary pathologies (Figure 1a). COX4I2 expression was elevated in the early stages of human COPD, defined by an FEV1 ≥ 80% pred., and in the murine model after 3 months of CS exposure (Figure 1b, c; Extended Data Table 1). However, in the later stages of COPD (FEV1 < 30% pred.) and after 8 months of CS exposure, the COX4I2 levels did not differ from those of the respective controls (Figure 1d, e; Extended Data Table 1). To investigate the role of COX4I2 in nitrosative stress, we used *ex vivo* murine lungs that were exposed to CS by intratracheal application in a repetitive manner (3 inhalations of 5 minutes each, with a one-hour interval between them) (Figure 1f). 3-Nitrotyrosine formation was largely prevented in isolated *Cox4i2^-/-^* lungs after CS exposure (Figure 1g, h).

**Figure 1.**
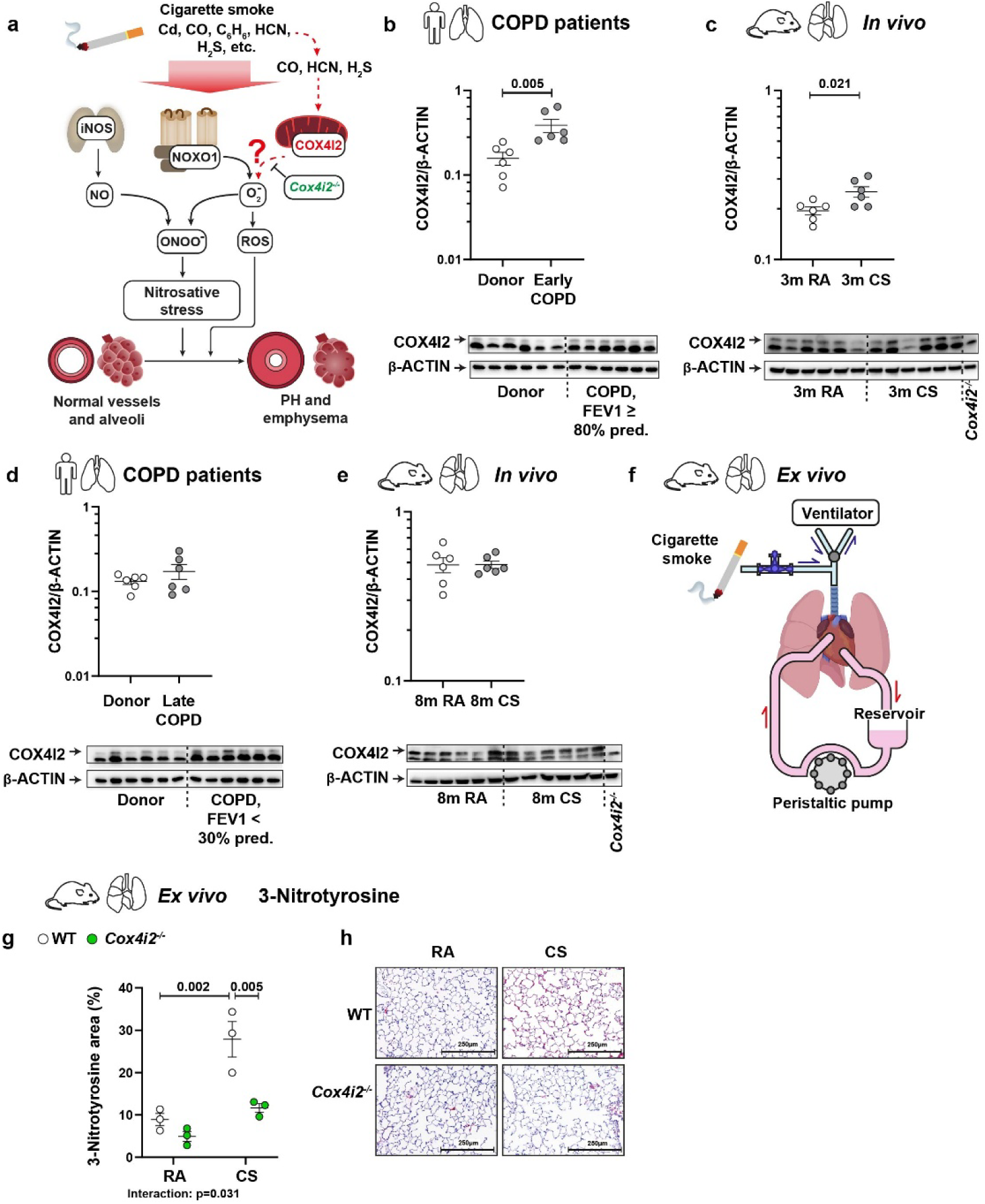
Upregulation of COX4I2 in early murine cigarette smoke-induced emphysema and human COPD and COX4I2-dependent nitrosative stress formation. a) Summary of the working hypothesis. Cigarette smoke (CS) exposure induces an increase in iNOS-generated nitric oxide (NO), which reacts with superoxide (O^2•–^) originating from NADPH oxidase and potentially mitochondria to form peroxynitrite (ONOO^-^). Peroxynitrite and potentially reactive oxygen species (ROS) induce the development of CS-induced emphysema and PH. b-e) Western blot analyses of COX4I2 protein expression in lungs explanted from healthy donors and people with early (b) and advanced stages (d) of COPD and from WT mice after 3 months (c) and 8 months (e) of CS or room air (RA) exposure. The specific signal from COX4I2 is marked by an arrow. Lung homogenate from *Cox4i2*^−/−^ mice was used as negative control. *n*=6 per group. f-h) Repeated application of CS in isolated lungs via tracheal application (3 times) for a period of 5 minutes once per hour (f) increased 3-nitrotyrosine formation in WT but not *Cox4i2^-/-^* lungs, as determined by 3-nitrotyrosine staining. g: Quantification of 3-nitrotyrosine staining. h: Representative images. *n*=3 per group. Statistical analysis was performed using Student’s t test and two-way ANOVA. Data from panels b-e were log-transformed prior to statistical analysis. The data are presented as the mean ± SEM. **Abbreviations**: Cd – cadmium; CO – carbon monoxide; C_6_H_6_ – benzene; HCN – hydrogen cyanide; H_2_S – hydrogen sulphide.

### *Cox4i2* deficiency protects against CS-induced emphysema but not PH

To assess the effect of *Cox4i2^-/-^*on the development of CS-induced emphysema and PH, we randomly assigned WT and *Cox4i2^-/-^* mice both genders to the CS-exposed group for 3 or 8 months (Extended Data Fig. 1a, Figure 2a). After 3 months of CS exposure, *Cox4i2^-/-^* mice exhibited fewer alterations of different lung functional parameters than WT mice did (Extended Data Fig. 1b-e). After 8 months of CS exposure, *Cox4i2^-/-^* mice were protected against CS-induced emphysema development, as demonstrated by *in vivo* lung functional assessments (Figure 2b, c, Extended Data Fig. 1f, g), µCT imaging (Figure 2d, Extended Data Fig. 1h, i), histological (Figure 2e, Extended Data Fig. 1j) and stereological analyses (Figure 2f, g, Extended Data Fig. 1k). In contrast, *Cox4i2^-/-^* mice exhibited PH after 3 months or 8 months of CS exposure to a similar degree as WT mice, as determined by *in vivo* haemodynamic measurements (Figure 2h, Extended Data Fig. 2a-c) and histological analysis of pulmonary vessels (Figure 2i, j). These findings suggest a specific protective effect of COX4I2 on alveolar destruction but not on vascular remodelling. Interestingly, *Cox4i2^-/-^* mice displayed significant protection against CS-induced alterations in the right ventricle (RV) only after 3 months but not after 8 months of CS exposure (Extended Data Fig 2d-i).

**Figure 2.**
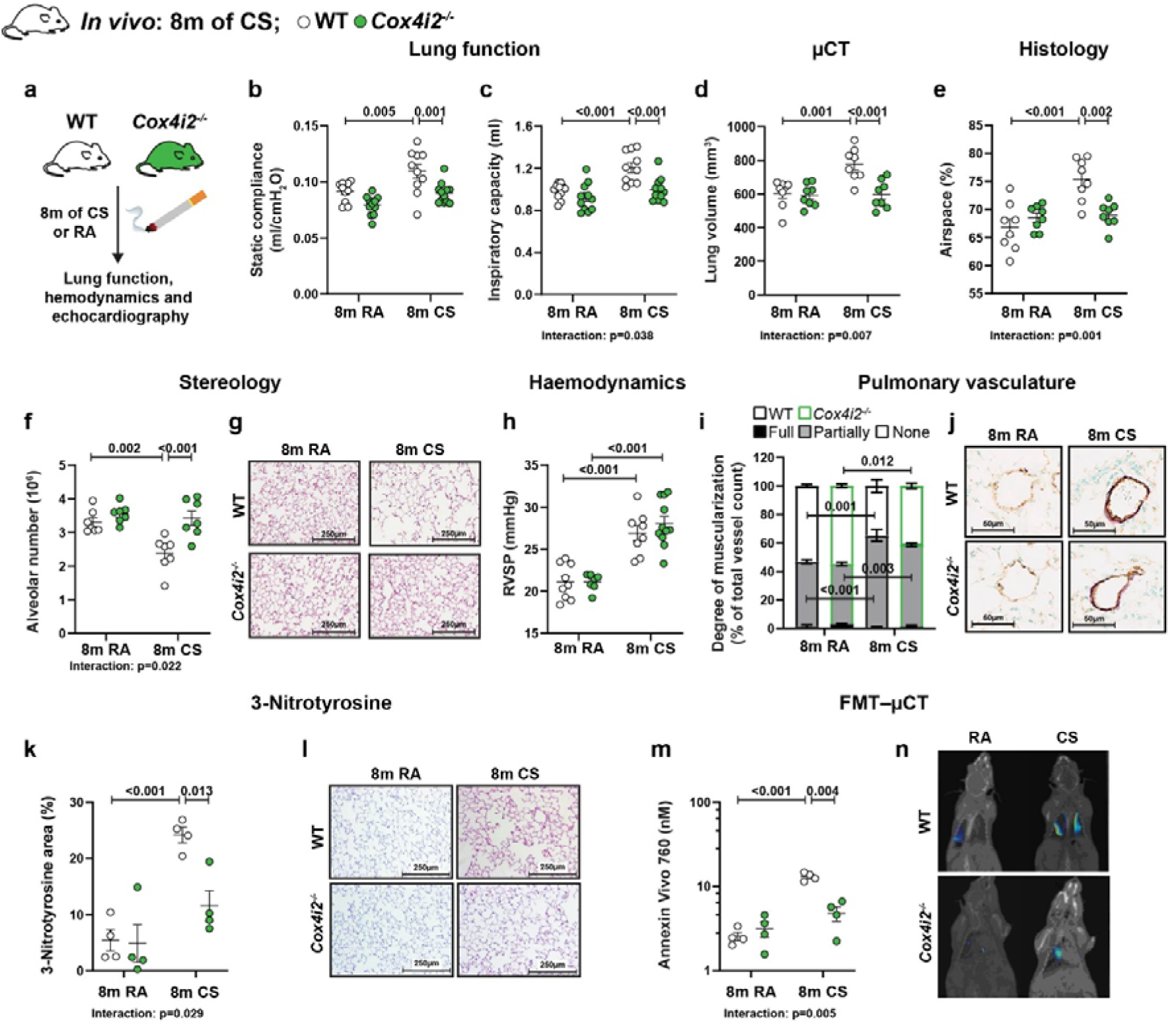
***Cox4i2* deficiency prevents the development of cigarette smoke-induced emphysema, apoptosis, and nitrosative stress but not pulmonary hypertension (PH).** a) WT and *Cox4i2^-/-^* mice were exposed to either cigarette smoke (CS) or room air (RA) for 8 months. b, c) Lung function after CS or RA exposure (*n*=10–12 per group): static compliance (b) and inspiratory capacity (c). d) µCT after CS or RA exposure (*n*=8 per group): Lung volume. e) Histological analysis of the lung parenchyma (*n*=8 per group): Airspace. f, g) Stereological analysis of the lung parenchyma (n=7 per group): alveolar number. g), Representative images of lung sections stained with haematoxylin and eosin (H&E). h) Haemodynamic measurements (*n*=7–11 per group): Right ventricular systolic pressure (RVSP). i, j) Morphological analysis of pulmonary vessels (*n*=4 per group): Data are provided for fully, partially, or nonmuscularized vessels as a percentage of the total vessel count. i: Quantification. j: Representative images. k, l) 3-Nitrotyrosine staining (*n*=4 per group). k: Quantification of 3-nitrotyrosine staining. l: Representative images. m, n) *In vivo* FMT-µCT measurements for quantification of apoptosis after CS or RA exposure (*n*=4 per group). m: Quantification. n: Representative images. Statistical analysis was performed using two-way ANOVA. Data from panel m were log-transformed prior to statistical analysis. The data are presented as the mean ± SEM.

In accordance with the protective effects of *Cox4i2^-/-^* on emphysema development, 3-nitrotyrosine staining was reduced in *Cox4i2^-/-^* mice compared with WT controls after 8 months of CS exposure (Figure 2k, l). Importantly, expression of *in vivo* apoptotic marker was also largely abolished in 8-month CS-exposed *Cox4i2^-/-^* mice compared with WT mice, as measured by fluorescence molecular tomography coupled with µCT imaging (Figure 2m, n).

### *Cox4i2* deficiency protects against CS-induced pulmonary inflammation

Inflammation is a central component of the pathogenesis of COPD^4^; however, the mouse model displays only mild inflammation. Nevertheless, the numbers of neutrophils, eosinophils, and T cells, but not macrophages, were highly elevated in bronchoalveolar lavage (BAL) samples from mice exposed to CS for 3 and 8 months, as determined by FACS analysis (Extended Data Fig. 3a-h, Figure 3a-d). *Cox4i2^-/-^* attenuated the increase in the numbers of neutrophils and T cells but not eosinophils or macrophages after 8 months but not after 3 months of CS exposure (Extended Data Fig. 3a-h, Figure 3a-d). Although *Cox4i2^-/-^*mice presented higher basal levels of CD4□ and CD8□ T cells than WT mice, after CS exposure, the number of T cells did not increase and was lower in *Cox4i2^-/-^* mice than in WT mice. Importantly, the infiltration of CD45^+^ cells (leukocytes) and CD3^+^ cells (T cells) after 8 months of CS was completely inhibited in *Cox4i2^-/-^* lung parenchyma (Figure 3e-h).

**Figure 3.**
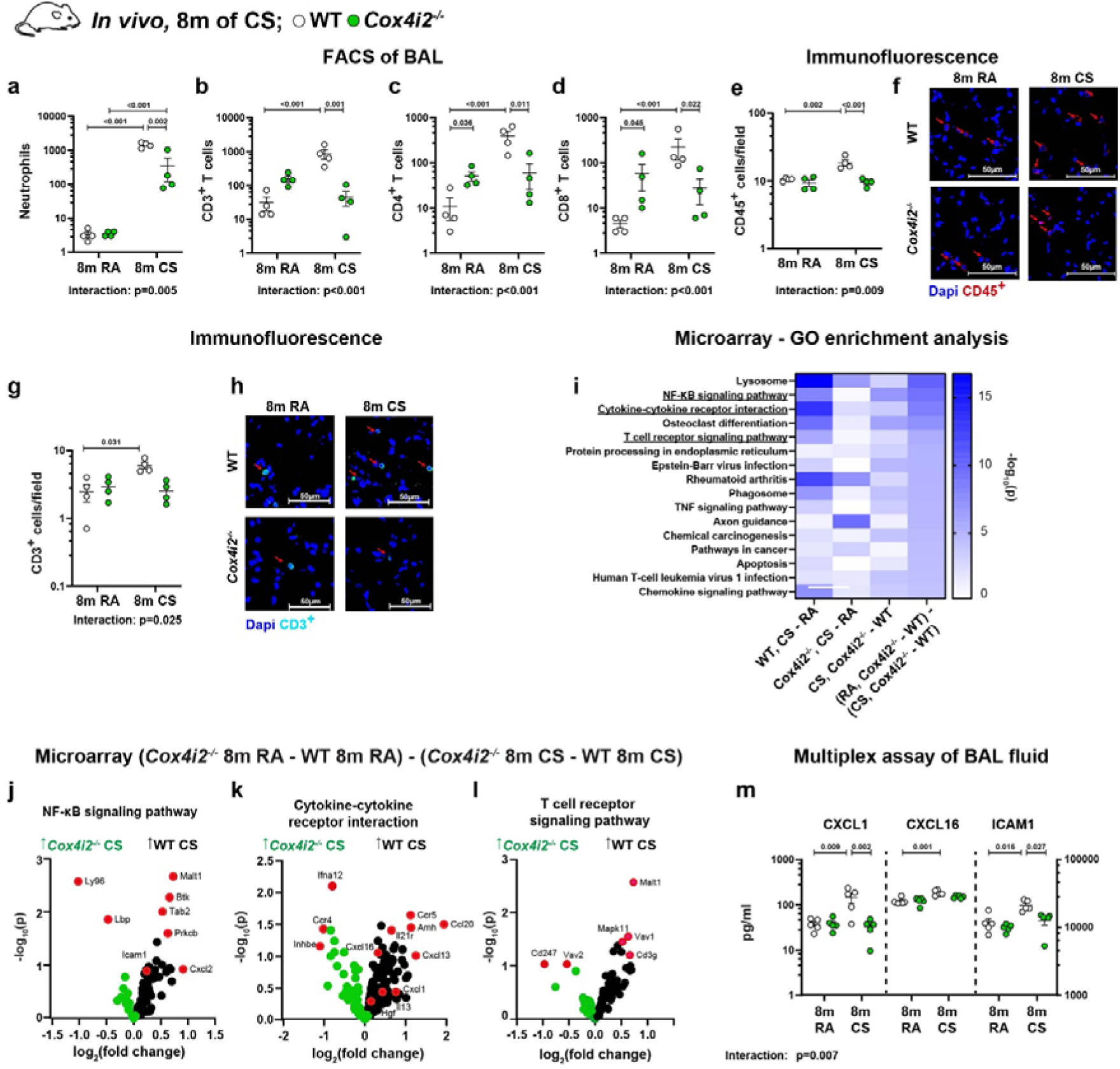
***Cox4i2* deficiency reduces inflammation in mice after 8 months of cigarette smoke exposure.** a-d) FACS analysis of bronchoalveolar lavage (BAL) from mice exposed to cigarette smoke (CS) or room air (RA) for 8 months (*n*=4 per group): Numbers of neutrophils (a), CD3^+^ T cells (b), CD4^+^ T cells (c) and CD8^+^ T cells (d). e-h) Immunofluorescence analysis of CD45^+^ (e, f) and CD3^+^ (g, h) cell accumulation in the lung parenchyma of mice exposed to CS or RA for 8 months (*n*=4 per group). e, g: Quantification. f, h: Representative images. i-l) Microarray analysis of lung homogenates from mice exposed to CS or RA for 8 months (*n*=6 per group). Most differentially regulated pathways between *Cox4i2*^−/−^ and WT mice (i). Differentially expressed genes involved in inflammatory pathways: NF-κB signalling (j), cytokine–cytokine receptor interaction (k) and T-cell receptor signalling (l). m) Multiplex analysis of BAL fluid (BALF) from mice exposed to CS or RA for 8 months (*n*=5 per group). Statistical analysis was performed using two-way ANOVA. Data from panels a-e, g and m were log-transformed prior to statistical analysis. The data are presented as the mean ± SEM.

Transcriptomic analysis of mRNA isolated from lung homogenates of mice exposed to CS for 8 months revealed differential regulation of inflammatory pathways in *Cox4i2^-/-^* lungs after CS (Figure 3i). Specifically, genes involved in NF-κB signalling, cytokine–cytokine receptor interactions, and T-cell receptor signalling were expressed at lower levels in *Cox4i2^-/-^* mice after CS exposure than in WT mice after CS exposure (Figure 3j-l). The levels of several proinflammatory chemokines/cytokines, such as CXCL1, CXCL16, and ICAM1, increased after CS exposure but were lower in CS-exposed BAL fluid (BALF) from *Cox4i2^-/-^* lungs than in WT lungs (Figure 3m). Accordingly, the levels of CXCL16 and ICAM1 were increased in the plasma of COPD patients (Extended Data Fig. 3i, j).

### *Cox4i2* is highly expressed in distinct pericyte subtypes

As COX4I2 expression was upregulated at early time points of CS exposure, we performed single-cell RNA sequencing (scRNA-Seq) of CD45^-^ and CD45^+^ lung cell suspensions from mice exposed to CS for 3 months to investigate the early triggers of COX4I2-mediated effects in different cell types (Figure 4a). A total of 10 distinct cell clusters were identified within the CD45^-^ population, and 16 distinct cell clusters in the CD45^+^ population (Figure 4b, Extended Data Fig. 4a-c, 5a, b, 6a-f). After CS exposure, ATI and ATII cells decreased, whereas all other cell types increased, with a particularly prominent increase in mesothelial cells, endothelial (EC), and alveolar capillary (aCap) cells in *Cox4i2^-/-^*lungs (Extended Data Fig. 4b; Extended Data Table 2). Among CD45^+^ cells, the most prominent changes after CS exposure were an increase in B cells and decrease in alveolar macrophages, proliferating T cells, and plasma cells (Extended Data Fig. 4c; Extended Data Table 2). Specifically, in *Cox4i2^-/-^* CS-exposed mice, the number of T cells increased, and the number of Treml4^+^ and Cd101^+^ neutrophils decreased although the latter showed higher levels in RA-exposed *Cox4i2^-/-^*compared to RA-exposed WT mice. In contrast, the number of Ngp^+^ neutrophils, which express markers of immature, recently recruited neutrophils from the bone marrow (Ly6c2^+^, Ly6G^+^, and Ngp^+^)^19^, was generally lower in *Cox4i2*^-/-^ mice and did not increase after CS exposure, unlike the increase observed in WT mice (Extended Data Fig. 4c; Extended Data Table 2). These data indicate a specific effect of *Cox4i2* deficiency on neutrophil regulation during CS exposure.

**Figure 4.**
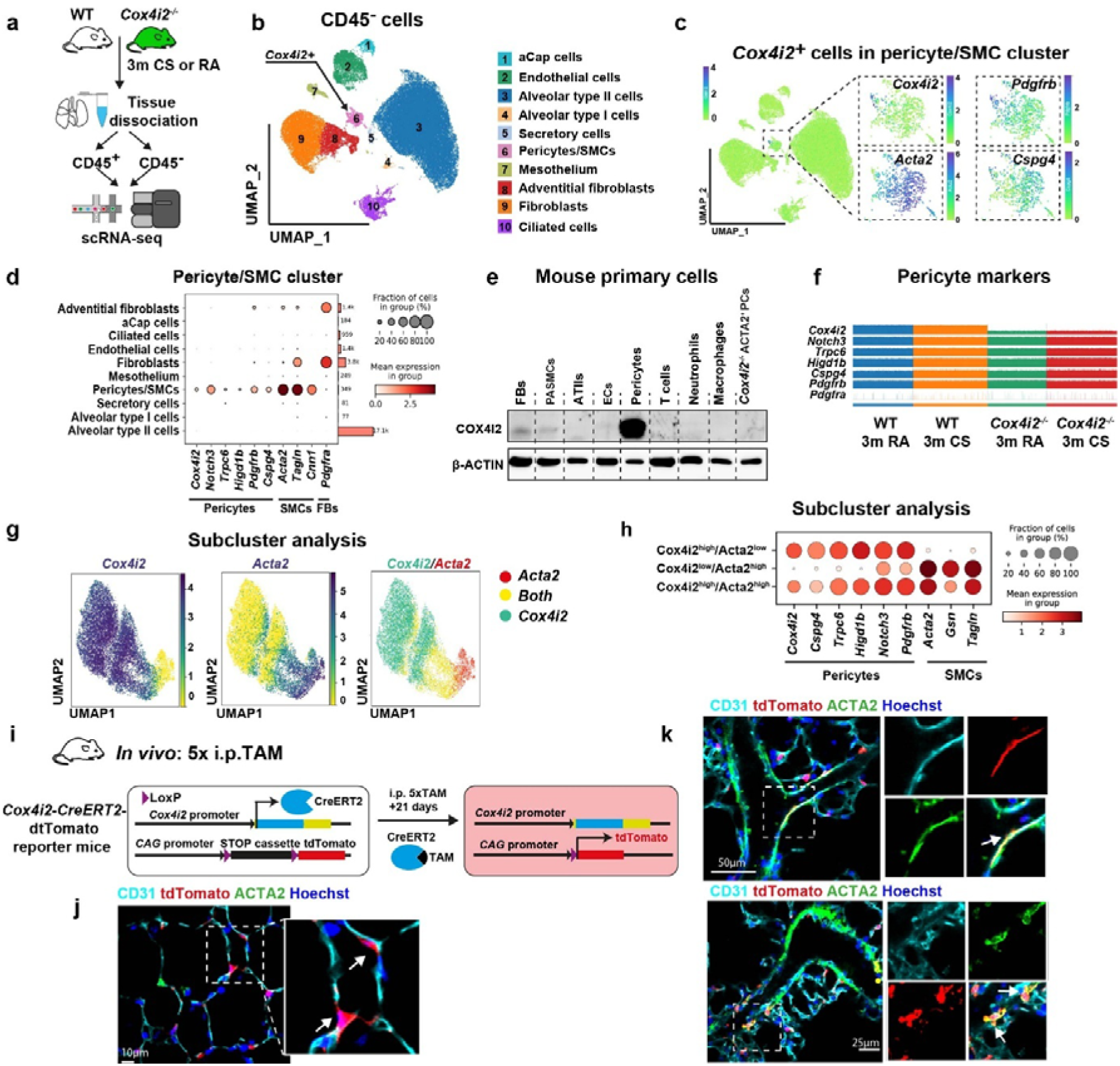
***Cox4i2* is predominantly expressed in pericytes, smooth muscle cells and fibroblasts** a) Schematic illustration of sample acquisition and data generation from CD45^+^ and CD45^-^ cells for single-cell RNAseq analyses. b) UMAP representation of CD45^-^ cells (*n*=2–3 per group). c) UMAP highlighting *Cox4i2* expressing cells in the pericyte/ smooth muscle cells (SMC) cluster of CD45^-^ cells. d) DotPlot visualization of marker genes for pericytes, SMCs and fibroblasts (FBs). e) Protein expression of COX4I2 in primary lung FBs, pulmonary artery SMCs (PASMCs), alveolar type II cells (ATIIs), endothelial cells (ECs), pericyte pellet, T cells, neutrophils and macrophages. PASMCs isolated from *Cox4i2*^−/−^ mice served as a negative control. f) Track plot of pericyte markers and fibroblast markers g) UMAP representation of the expression of *Cox4i2*, *Acta2*, or both in pericytes. Red: cells expressing *Acta2*; green: cells expressing *Cox4i2*; yellow: cells expressing both *Acta2* and *Cox4i2*. h) DotPlot visualization of marker genes expressed in pericytes. i) Labelling of *Cox4i2*-expressing cells was performed using *Cox4i2*-CreERT2-tdTomato mice. A Kozak-CreERT2-P2A cassette was inserted just before the *Cox4i2* start codon, and these mice were crossed with tdTomato reporter mice containing a loxP-STOP-loxP cassette. After tamoxifen (i.p.; TAM) injection, Cre activation removed the stop cassette, enabling tdTomato expression driven by the *Cox4i2* promoter. j, k) Representative images. j: tdTomato-positive cells in alveolar structures costained with CD31 as an endothelial marker and ACTA2 as an SMC marker. k: tdTomato-positive cells in pulmonary vessels. Colour code: Red – tdTomato, green – ACTA2, bright cyan – CD31, blue – Hoechst.

High *Cox4i2* transcript expression was detected in a cluster composed of cells expressing pericyte and SMC markers (Figure 4b-d, Extended Data Fig. 6g), named the pericyte/SMC cluster. Moreover, analysis of the integrated Human Lung Cell Atlas (HLCA), which combines 49 datasets of the human respiratory system into a single atlas encompassing more than 2.4 million cells from 486 individuals^20^, revealed that *COX4I2* is highly expressed in pericytes and to a lesser extent in SMCs (Extended Figure 7a). We validated the scRNA-Seq findings by Western blot analysis and detected COX4I2 protein expression in pericytes, precapillary PASMCs and lung fibroblasts (FBs) (Figure 4e). To understand the cell type-specific effects of *Cox4i2^-/-^* on different signalling pathways after CS, we performed gene set enrichment analysis (GSEA) on hallmark gene sets from the Molecular Signatures Database (MSigDB). Our analysis revealed that among CD45^-^ cells, the *Cox4i2^+^*cell containing pericyte/SMC cluster showed differential regulation of various pathways, including oxidative phosphorylation and *Tnf*_α_, in CS-exposed *Cox4i2^-/-^* mice compared with those in WT mice (Extended Data Fig 7b). With respect to CD45^+^ cells, we focused on neutrophils, as neutrophil migration is regulated by pericytes^21^ and their numbers were lower in the BAL of *Cox4i2^-/-^* CS mice. Ngp□ neutrophils, whose abundance was specifically downregulated in *Cox4i2^-/-^* CS mice compared with that in WT CS mice (Extended Data Table 2), exhibited the strongest transcriptional response to CS exposure (Extended Fig. 7c-e). To gain deeper insights into the cell population that highly expresses *Cox4i2*, we established a modified^22^ cell isolation protocol using MACS- or FACS-based negative selection for CD45 (immune cells), CD31 (ECs), CD326 (epithelial cells) and CD140a (PDGFRA), followed by positive selection for CD140b (PDGFRB). Importantly, *Cox4i2* expression decreased after passaging; thus, we used only cells at passage zero for further *in vitro* experiments (Extended Data Fig. 8a). Using FACS-based cell sorting, we performed scRNA-seq on pericytes isolated after 3 months of exposure to CS or RA (Extended Data Fig. 8b). We confirmed that the isolated cells were pericytes as they expressed specific pericyte markers, such as *Notch3, Trpc6*, *Higd1b, Pdgfrb,* and *Cspg4*, and did not express the FB marker *Pdgfra* (Figure 4f). Pericytes exhibited increased mtROS levels from WT but not *Cox4i2^-/-^* lungs after 3 m of CS exposure (Extended Data Fig. 8c, d). *Acta2* was expressed only in a specific cell subcluster and only partially overlapped with *Cox4i2* expression (Figure 4g, h). Therefore, we defined three subclusters similar to those proposed previously^23^: Cox4i2^high^/Acta^low^ cells (“pericytes supporting EC” or “classical pericytes”), Cox4i2^high^/Acta2^high^ cells (“transitional pericytes”) and *Cox4i2*^low^/*Acta2*^high^ (“contractile pericytes”). GO analysis revealed the most prominent genotype-specific difference in gene regulation in the “classical pericyte” Cox4i2^high^/Acta2^low^ subcluster (Extended Data Fig. 8e-g).

To identify different *Cox4i2*-expressing cell populations in the lung, we generated transgenic mice in which the fluorescent protein tdTomato was expressed under the control of the *Cox4i2* promoter (*Cox4i2*-CreERT2-tdTomato) following intraperitoneal (i.p.) administration of tamoxifen (TAM) (Figure 4i). Using these mice, we detected *Cox4i2* expression mostly in the lung parenchyma close to capillaries (Figure 4j, Extended Data Video 1) and rarely within the vascular precapillary ACTA2^+^ cells (Figure 4k), confirming the existence of two distinct *Cox4i2*^+^ cell populations with low or high levels of ACTA2 expression. This latter population is likely captured within the isolated precapillary PASMCs. The control mice did not express any tdTomato in the lung (Extended Data Figure 9a). Additionally, we localized distinct *COX4I2*-expressing cell populations in the human lung using a combination of *in situ* hybridization to detect *COX4I2* mRNA and immunofluorescence staining for specific cell markers (SMCs: ACTA2; ECs: von Willebrand factor; pericytes: PDGFRB/CSPG4). Consistent with observations in mouse lungs, *COX4I2* expression in human lungs was primarily detected in PDGFRB-positive cells and rarely detected in *ACTA2*-positive cells (Extended Data Figure 9b).

### *Cox4i2* expressing pericytes promote neutrophilic inflammation and angiogenesis

To understand the protective effects of *Cox4i2^-/-^* on inflammation, which we detected in CS-exposed *Cox4i2^-/-^* mice *in vivo*, and as we detected anti-inflammatory effects in Ngp^+^ neutrophils, we investigated neutrophilic inflammation in pericytes *in vitro*. We measured neutrophil migration *in vitro* after the application of media from WT or *Cox4i2^-/-^* pericytes that had been stimulated with the TLR4 agonist lipopolysaccharide from *Escherichia coli* 0111: B4 (LPS-EB). Neutrophilic migration was enhanced in the presence of supernatant from stimulated pericytes; however, this effect was weaker in the *Cox4i2^-/-^* supernatant than in the WT supernatant (Figure 5a). The inhibition of CXCL1 activity, which plays a crucial role in recruiting neutrophils^24^, by 10 µM SX-517 inhibited LPS-EB-induced migration in WT mice but did not further decrease migration in *Cox4i2^-/-^*mice (Figure 5a). Furthermore, neutrophil migration was inhibited by treatment with 10□nM of the mitochondria-targeted antioxidant MitoQ^25^, (Figure 5b). Similarly, LPS-EB-induced mtROS production was lower in *Cox4i2^-/-^* pericytes than in WT pericytes (Extended Data Fig. 10a, b). Additionally, we tested the effect of a selective inhibitor of mtROS release from complex III^26^, S3QEL, on neutrophil migration. S3QEL inhibited LPS-EB-induced neutrophil migration in the supernatant of WT but not *Cox4i2^-/-^*pericytes (Figure 5c). Accordingly, both the mRNA of *Cxcl1* in pericytes and its protein level in the cell culture supernatant were less elevated in *Cox4i2^-/-^* pericytes after TLR4 stimulation than in WT pericytes (Figure 5d-e), similar to the CXCL1 levels in the BAL fluid of 8 m CS-exposed mice (Figure 3m). Furthermore, MitoQ and S3QEL treatment inhibited both the mRNA expression of *Cxcl1* in pericytes and its protein level in the supernatant of WT pericytes stimulated with a TLR4 agonist (Figure 5f-g, Extended Data Fig. 10c, d). Although the TLR4-mediated upregulation of CCL2 and IL6 was not impaired in *Cox4i2^-/-^*pericytes, MitoQ still reduced their levels in the supernatant, suggesting that, in addition to COX4I2, other mitochondrial oxidative stress mechanisms may contribute to their regulation (Extended Data Fig. 10e-h).

**Figure 5.**
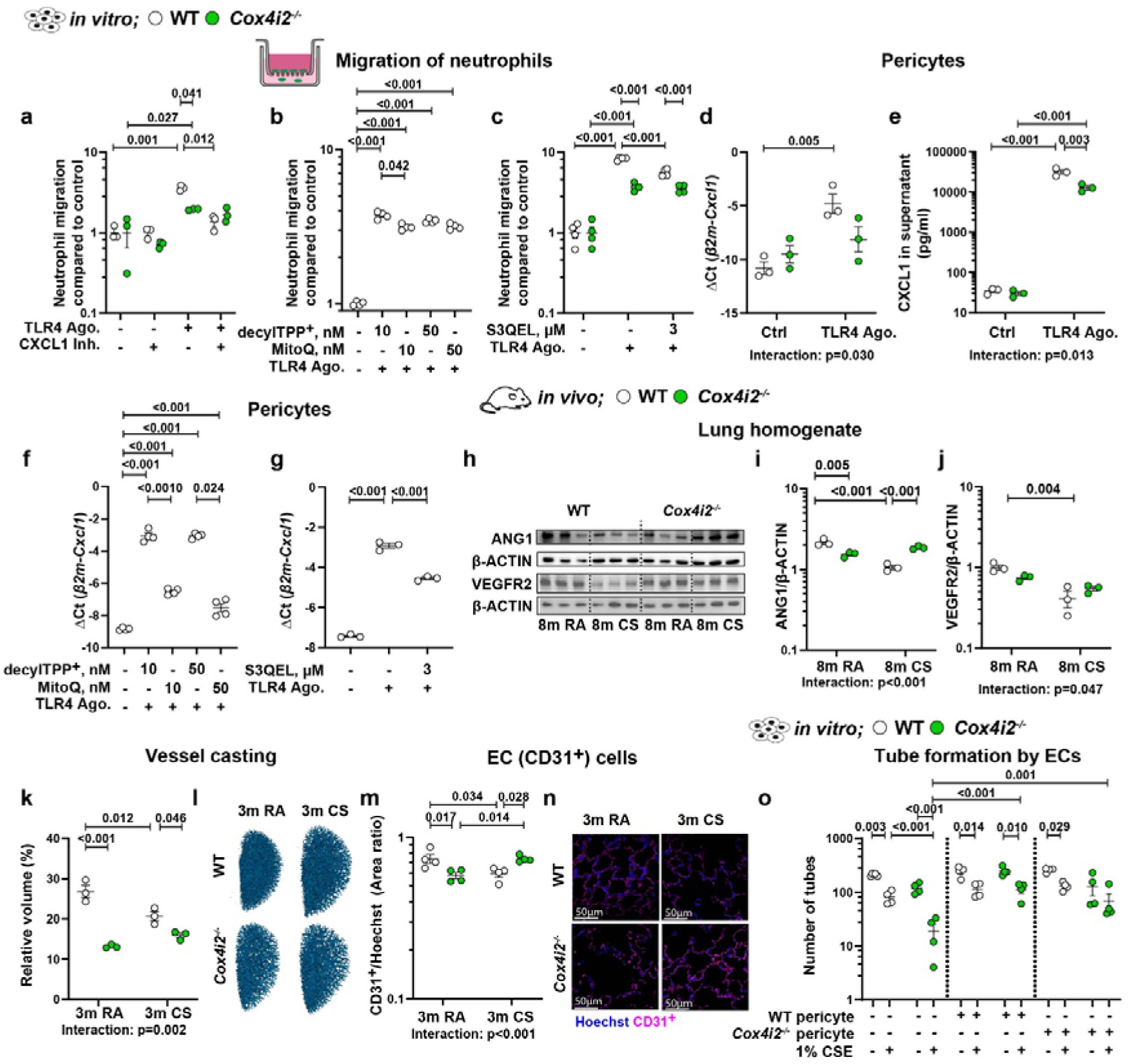
The effects of *Cox4i2* deficiency on neutrophilic inflammation and angiogenesis. a-c) Migration of neutrophils in response to conditioned medium from pericytes isolated from *Cox4i2*^−/−^ and WT mice, following stimulation with the TLR4 agonist LPS-EB and treatment with the CXCL1 inhibitor 10µM SX-517 (a, *n*=3) and from WT mice treated with different doses of MitoQ/DecylTPP^+^ (b, *n*=4) or S3QEL (c, *n*=4). d, e) mRNA expression of *Cxcl1* in pericytes isolated from *Cox4i2*^−/−^ and WT mice and exposed to the TLR4 agonist LPS-EB (d, *n*=3) or CXCL1 protein in the cell culture supernatant (e, n=3). f, g) mRNA expression of *Cxcl1* in pericytes isolated from WT mice and stimulated with the TLR4 agonist LPS-EB following treatment with MitoQ/DecylTPPLJ (f, *n* = 4) or S3QEL (g, *n*=3). h-j) Protein expression of ANG1 (i) and VEGFR2 (j) in the lung tissues of *Cox4i2*^−/−^ and WT mice exposed to CS for 8 months. *n*= 3 per group. h: Representative images. k, l) Pulmonary vessel casting of mice exposed to CS for 3 months (*n*=3 per group). l: Representative images. m, n) Number of CD31^+^ cells (ECs) in the lung parenchyma of *Cox4i2*^−/−^ and WT mice exposed to CS for 3 months (m, *n*= 4 per group). n: Representative images. o) Vascular tube formation of ECs incubated with solvent or cigarette smoke extract (CSE) in the presence or absence of primary pulmonary pericytes (*n*= 4, number of tubes). Statistical analysis was performed using one-way and two-way ANOVA. Data from panels a-c, e, i, j, m and o were log-transformed prior to statistical analysis. *n* in figures a-g and o represents individual cell isolations per group. The data are presented as the mean ± SEM.

Given that pericytes play a central role in angiogenesis^27^, and as we found effects of *Cox4i2* deficiency on EC number, we further investigated the effect of *Cox4i2* deficiency on angiogenesis. Indeed, the expression of angiopoietin 1 (ANG1), which is important in the pericytes for vessel maturation^28^, was significantly greater in lungs of 8 m CS *Cox4i2^-/-^* mice than in 8 m CS WT mice. However, ANG1 expression was lower in RA-exposed *Cox4i2^-/-^*mice than in WT mice (Figure 5h, i). VEGFR2 expression, which controls angiogenesis^29^, tended to show similar regulation as ANG1 expression (Figure 5h, j). Accordingly, vessel casting revealed a lower vascular volume in *Cox4i2^-/-^* lungs than in WT lungs during both RA and CS, although the vascular volume was unchanged after 3 m of CS exposure in these mice but decreased in WT mice (Figure 5k, l). Interestingly, compared with WT lungs, *Cox4i2^-/-^*lungs exhibited fewer CD31^+^ ECs at 3 m RA but more ECs after 3 m CS (Figure 5m, n).

The viability of pericytes isolated from *Cox4i2^-/-^* mice was similar to that of pericytes isolated from WT mice in response to CS extract (CSE) *in vitro* (Extended Data Fig. 10i), and at these doses of CSE mtROS levels were similar in both genotypes (Extended Data Fig. 10j), suggesting a paracrine effect of pericytes on angiogenesis. In accordance with vessel casting, *Cox4i2^-/-^* ECs formed fewer vascular tubes when they were cocultured with *Cox4i2^-/-^* pericytes in RA compared with WT ECs (Figure 5o, Extended Data Fig. 10k). Furthermore, *in vitro* CSE exposure reduced vascular tube formation in ECs isolated from WT and *Cox4i2^-/-^* mice, but this effect was more severe in *Cox4i2^-/-^* mice. Coculture with pericytes from WT or *Cox4i2^-/-^*mice attenuated the latter effect. Additionally, we examined the effect of *Cox4i2^-/-^*on the retinal capillary plexus, as it serves as an established model system for studying angiogenesis^30^. Consistent with the lung findings, the capillary plexus was denser in WT mice than in *Cox4i2^-/-^* mice in the control nonsmoking group. However, after 3 months of CS exposure, the density of the capillary plexus in WT mice was lower than that in *Cox4i2^-/-^* mice (Extended Data Fig. 11). These results suggest that *Cox4i2* deficiency inhibits angiogenesis during development but may (indirectly) support ECs during *in vivo* CS exposure via systemic effects.

As COX4I2 is also expressed in precapillary PASMCs (Figure 4e), which partially resemble Cox4i2^low^/Acta2^high^ and Cox4i2^high^/Acta2^high^ cells, we investigated the functional relevance of *Cox4i2* deficiency in precapillary PASMCs on CSE-induced effects *in vitro*. Exposure to 3% CSE did not affect the viability of WT or precapillary *Cox4i2^-/-^* PASMCs (Extended Data Fig. 12a). We thus used 3% CSE for further experiments. Precapillary *Cox4i2^-/-^* PASMCs were protected against CSE-induced mitochondrial oxidative stress (Extended Data Fig. 12b, c) and CSE-induced necrosis (Extended Data Fig. 12d, e). Interestingly, compared with medium from precapillary WT PASMCs, conditioned medium from precapillary *Cox4i2^-/-^*PASMCs protected ATII cells from CSE-induced reductions in cell viability (Extended Data Fig. 12f). These findings highlight the potential paracrine effect of *Cox4i2* deficiency in mitigating CSE-induced alveolar epithelial cell damage.

To demonstrate the deleterious effect of COX4I2 on CSE-induced cellular responses, we overexpressed *Cox4i2* in the murine alveogenic lung carcinoma cell line CMT64/61, which does not naturally express *Cox4i2* (Extended Data Fig. 13a). The overexpression of *Cox4i2* exacerbated the harmful effects of CSE on cell viability (Extended Data Fig. 13b). Additionally, *Cox4i2* overexpression significantly increased mitochondrial oxidative stress levels at baseline and after CSE exposure (Extended Data Fig. 13c, d). While *Cox4i2* overexpression did not alter basal necrosis levels, it enhanced CSE-induced necrosis (Extended Data Fig. 13e).

### MitoQ treatment reverses inflammation, emphysema and PH after 8 months of CS exposure

To investigate whether inhibition of mtROS generation and/or oxidative stress could be a therapeutic strategy for the treatment of COPD, we treated mice with the mitochondria-targeted antioxidant MitoQ (50 mg/kg/day) for 3 months after 8 months of exposure to CS. As shown previously, this protocol induces irreversible emphysema and PH in untreated mice^8,9^. MitoQ acts against peroxynitrite and mitochondrial lipid peroxidation. As a control, mice were treated with the inactive carrier substance decylTPP^+^ (Figure 6a). After 3 months of MitoQ treatment subsequent to 8 months CS exposure, the CS-induced changes in lung function were attenuated (Figure 6b, Extended Data Fig. 14a, b), and the number of alveoli was restored (Figure 6c, d; Extended Data Fig. 14c). Moreover, MitoQ treatment normalized right ventricular systolic pressure (RVSP) (Figure 6e, Extended Data Fig. 14d) and restored pulmonary vasculature remodelling (Figure 6f, g) and RV dilatation and function (Extended Data Fig. 14e-g). Importantly, MitoQ treatment significantly reduced the accumulation of CD45^+^ leukocytes and CD3^+^ T cells within the septal walls of the lung parenchyma (Figure 6h, i, Extended Data Fig 14h, i) and decreased nitrosative stress, as indicated by decreased 3-nitrotyrosine accumulation (Figure 6j, k). Consistently, treatment with 50 nM MitoQ mitigated the effect of 5% CSE on cell proliferation in mouse precision-cut lung slices (PCLS) (Figure 6l, m). Furthermore, MitoQ treatment attenuated the CS-induced increase in CXCL1 and CXCL16 levels in the BALF (Extended Data Fig. 14j), similar to the effect observed in *Cox4i2^-/-^* mice (Figure 3m).

**Figure 6.**
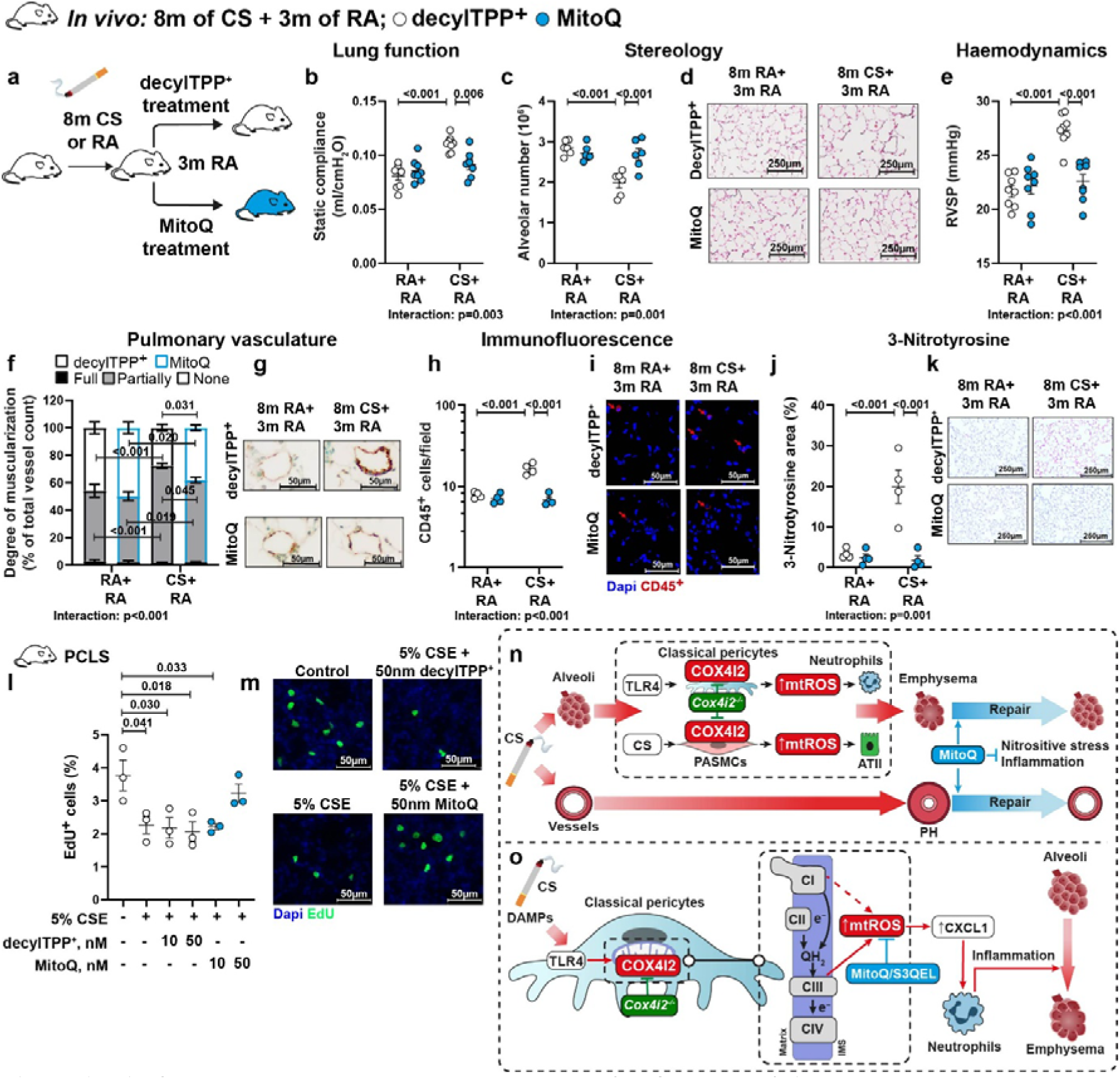
MitoQ treatment reverses emphysema and PH after 8 months of cigarette smoke exposure. a) WT mice were exposed to room air (RA) or cigarette smoke (CS) for 8 months, followed by treatment with MitoQ (50 mg/kg/day) or the inactive carrier decylTPP+ for 3 months via gavage. b) Lung functional analyses (*n*=7-8 per group) of static compliance. c-d) Stereological analysis of the lung parenchyma (*n*=6 per group) showing the number of alveoli (c) and representative pictures (d). e) Haemodynamic measurements (*n*=8 per group): right ventricular systolic pressure (RVSP). f, g) Morphological analysis of pulmonary vessels (*n*=5 per group): Data are provided for fully, partially, or nonmuscularized vessels as a percentage of the total vessel count. f: Quantification. g. Representative images. h, i) Immunofluorescence analysis (*n*=4 per group) of CD45^+^ cell accumulation in the lung parenchyma. h: Quantification. i: Representative images. j, k) 3-Nitrotyrosine staining of lung tissue (*n*=4 per group). j: Quantification of nitrotyrosine staining. k: Representative images. l, m) Percentage of EdU-positive cells in mouse PCLS treated with 5% CSE in the presence of either MitoQ or DecylTPPLJ (*n*=3 per group). l: Quantification of EdU-positive cells. m: Representative images. Statistical analysis was performed using one-way and two-way ANOVA. Data from panels h were log-transformed prior to statistical analysis. The data are presented as the mean ± SEM. n, o) Summary of findings showing the critical role of COX4I2 in CS-induced pulmonary inflammation and emphysema development and the therapeutic potential of MitoQ (n) and COXI2-dependent regulation of neutrophil migration via pericytes (o).

## Discussion

Our study shows for the first time that COX4I2 is predominantly expressed in specific subtypes of pericytes and precapillary PASMCs and is essential for the regulation of pulmonary inflammation, apoptosis and microvascular homeostasis. COX4I2 deficiency in mice inhibits the development of CS-induced nitrosative stress, inflammation and pulmonary emphysema. With respect to cellular mechanisms, we revealed that increased COX4I2-dependent mtROS generation and/or oxidative stress induced by either TLR4 signalling in classical pericytes or direct CSE-mediated effects in precapillary PASMCs regulates neutrophil migration and ATII apoptosis, respectively. Accordingly, global protection against mitochondrial oxidative stress with MitoQ reverses pulmonary inflammation and emphysema. Furthermore, in contrast to the effect of knocking out of *Cox4i2*, MitoQ also reversed PH (Figure 6n, o). In this regard, loss of *Cox4i2* inhibits developmental angiogenesis and CS-induced microvascular alterations but does not impact PH, emphasizing the cell type-specific effects of COX4I2 and compartmental regulation of COPD pathologies.

Our findings build on our previous investigations that highlight the crucial role of oxidative and nitrosative stress in the development of COPD through the promotion of inflammation and apoptosis^8,9^. Moreover, a previous study suggested that the upregulation of COX4I2 expression may contribute to CS-induced alterations in COX function and subsequent mitochondrial dysfunction, which promotes CS-induced hyperinflammation^18^. Interestingly, in our study, COX4I2 was upregulated only during the early stages of murine CS-induced emphysema and human COPD, which highlights COX4I2-dependent mtROS production and mitochondrial oxidative stress and/or damage as potential triggers of disease development. Accordingly, *Cox4i2* deletion inhibited CS-induced inflammation and emphysema after 8 months of CS exposure, and the effects first occurred after 3 months of CS exposure. Importantly, *Cox4i2* deletion did not affect the development of PH. We previously showed that CS-induced PH development is dependent on iNOS activity in bone marrow-derived cells^10^, which is consistent with COX4I2 exerting cell type-specific effects. In this regard, our microarray analysis suggested that inflammatory pathways are affected, particularly immune cell infiltration in the lung. However, scRNA-seq revealed that *Cox4i2* was not expressed to a detectable level in immune cells but was expressed predominantly in cell populations that expressed SMC and/or pericyte markers. We identified a) “classical pericytes” (that is, “pericytes supporting ECs”) expressing typical pericyte markers (*Higd1b*, *Notch3*, *Cspg4,* and *Pdgfrb)* and low levels of SMC markers (e.g., *Acta2)*^23^, b) a subpopulation with features resembling pericytes and SMCs (Cox4i2^high^, Pdgfrb^high^, and Acta2^high^), which may be regarded as “transitional pericytes” (previously described as “oxygen-sensing PASMCs”^23^) and c) a subpopulation of pericytes resembling stronger features of SMCs (Cox4i2^low^/Acta2^high^,“contractile pericytes”). Using genetically modified mice expressing tdTomato under the *Cox4i2* promoter, we found “classic pericytes” (Cox4i2^high^, Acta2^low^) located in the capillary region of alveoli, representing the majority of tdTomato^+^ cells. In contrast, fewer cells are located in the precapillary transitional zone between muscularized and non-muscularized pulmonary microvessels and may represent PASMCs or “transitional” and “contractile” pericytes (precapillary ACTA2+ cells). Consistent with our findings, Klouda *et al*., who used *Higd1b* as a pericyte marker, identified two subtypes of lung pericytes: type 1, which supports capillary homeostasis, and type 2, which coexpresses SMC markers^31^. These contractile pericytes, in particular, can regulate the direction of blood flow at capillary junctions^32^, although whether they express COX4I2 is not known. Analysis of publicly available scRNA-seq data from human lung tissue revealed that, similar to that in mouse lungs, COX4I2 is predominantly expressed in pericytes and SMCs.

To understand the role of COX4I2 in pericytes, we focused on neutrophil migration and angiogenesis, as both are regulated by pericytes^33,34^, and our data indicated an effect of *Cox4i2* deficiency on specific neutrophils (Ngp^+^) and ECs. *Cox4i2^-/-^* suppressed CS-induced expression of CXCL1, a key driver of neutrophil recruitment^24^, *in vivo* and suppressed CXCL1expression and CXCL1-dependent neutrophil migration in TLR4-stimulated pericytes *in vitro*. We further demonstrated that COX4I2 regulates the expression and secretion of CXCL1 and neutrophil migration through complex III-induced oxidative stress, as TLR4-stimulated CXCL1 expression and neutrophil migration are inhibited by the mitochondria-targeted antioxidant MitoQ^25^ and by S3QEL, which has been shown to inhibit mtROS release specifically from complex III^26^. These findings are consistent with prior evidence that TLR4 activation increases mtROS release from complexes I and III in macrophages and T cells, thereby promoting the production of proinflammatory cytokines^16,17^. As, we did not detect a direct effect of low dose CSE on mtROS production in pericytes, we conclude that the effects on pericyte during CS exposure are transferred via TLR4-COX4I2-mtROS signalling but not direct effects of CS on COX4I2-induced mtROS release via interaction with ETC components.CXCL1 expression was previously shown to be upregulated in BAL fluid from COPD patients^35^ and thereby can attract neutrophils in COPD. CXCL1 signalling in turn can be activated by CS-dependent cellular injury and release of damage-associated molecular patterns (DAMPs)^36–38^. In COPD, activated neutrophils release proteolytic enzymes such as neutrophil elastase and matrix metalloproteinases (MMPs), leading to the degradation of extracellular matrix components, particularly elastin, and thus are key drivers of emphysema development^3,39^. Accordingly, we found that different *in vivo* neutrophil subpopulations were altered in *Cox4i2*^-/-^ CS mice, particularly Ngp□ neutrophils (immature, recently recruited neutrophils^19^), and numbers of CD45^+^ and CD3^+^ cells were decreased in the lung parenchyma after 8 months of CS exposure. In contrast to *Cox4i2^-/-^*, MitoQ also suppressed the secretion of CCL2 and IL-6, suggesting that MitoQ exerts effects beyond those mediated by blocking the action of COX4I2 alone. Interestingly, *Cox4i2^-/-^* inhibited the CS-induced increase in ICAM1 and CXCL16 in the BALF of mice exposed to CS for 8 months, whereas these cytokines were not detected or not regulated in the cell culture supernatant from pericytes following TLR4 stimulation. These findings suggest that these cytokines are either not regulated by TLR4 signalling or are not secreted by pericytes, and that other cell types may contribute to the protective effects of *Cox4i2* deficiency. In this context, it is notable that alveolar epithelial cells express ICAM1, which is required for neutrophil migration into the airways *in vivo*^40^. Moreover, we detected increased levels of ICAM1 and CXCL16 in human plasma from COPD patients, suggesting the involvement of similar mechanisms in human disease.

Notably, COX4I2 was also essential for vascular tube formation *in vitro* and vascular development *in vivo*. Although *Cox4i2* expression was not detected in ECs, *Cox4i2^-/-^* ECs exhibited an increased inhibitory effect of CSE on vascular tube formation. While the underlying mechanisms remain unclear, COX4I2 in pericytes may influence EC epigenetics. Although no literature directly confirms this hypothesis, pericytes and ECs from different organs exhibit distinct epigenetic landscapes^41^. Interestingly, compared with WT ECs, *Cox4i2^-/-^* ECs formed fewer vascular tubes when cocultured with *Cox4i2^-/-^* pericytes in RA, and only the coculture of *Cox4i2^-/-^* ECs with *Cox4i2^-/-^* pericytes, but not with WT pericytes, protected against the effects of CSE exposure on tube formation. Accordingly, although unstressed *Cox4i2^-/-^*mice exhibit a lower vascular volume, they lack a further CS-induced decrease, as was observed in WT mice. Given that quantifying the capillary network in the lung is challenging, we also investigated the retinal vasculature as a prototypical vascular network. Interestingly, we found decreased network formation in *Cox4i2*^-/-^ RA mice compared with WT RA mice but improved network formation after CS. Thus, *Cox4i2* deletion inhibits developmental angiogenesis but may have a protective effect on CS-induced vascular pruning, which is among the characteristics of COPD. Retinal effects after CS exposure may be transmitted via systemic effects of CS exposure which can induce retinal inflammation, as we demonstrated previously^42^, and these effects may be inhibited in *Cox4i2*^-/-^ mice. Importantly, pulmonary arterial pruning has been associated with faster progression of emphysema and a more rapid decline in lung function in human studies^43^. Although the role of pericytes in COPD development is poorly understood, our findings agree with those of previous studies that revealed impaired pericyte migration in *in vivo* murine emphysema models^44^ and decreased pericyte-EC interdigitations in the lungs of COPD patients^45^. Moreover, it is well-established that CS is an independent risk factor for decreased vessel density in the deep retinal capillary plexus^46^.

In addition to pericytes, classically isolated precapillary PASMCs, which also express high levels of COX4I2 and may represent a mixture of true precapillary PASMCs and pericyte subpopulations such as “transitional pericytes” and “contractile pericytes”, here collectively termed precapillary ACTA2□ cells, may contribute to COX4I2-dependent effects. As shown for acute hypoxia^14^, CSE-exposed *Cox4i2*^-/-^ PASMCs generated fewer mtROS, and conditioned medium from *Cox4i2^-/-^* PASMCs protected ATIIs from CSE-induced necrosis. Notably, PASMCs from COPD patients exhibit accelerated senescence and increased release of soluble and insoluble paracrine factors^47^, which may contribute to the detrimental effects of CS on ATII cells. The critical role of COX4I2-dependent mtROS generation and/or oxidative stress is further supported by our findings in *Cox4i2*-overexpressing CMT cells, which exhibit increased CSE-induced oxidative stress and necrosis, along with decreased viability. The molecular mechanism through which the expression of COX4I2 in the COX complex alters mtROS production from the ETC and oxidative stress remains unknown and will be addressed in the future.

In conclusion, COX4I2 is a novel key factor in distinct pericyte subtypes and in the development of CS-induced inflammation and emphysema by regulating neutrophil migration, angiogenesis and apoptosis. Cell type-specific effects contribute to *Cox4i2^-/-^*-mediated protection specifically against CS-induced emphysema, but not PH. In contrast, global protection against mitochondrial oxidative stress by MitoQ inhibited inflammation, emphysema and PH, notably even when treatment started after disease establishment, and thus may offer a novel therapeutic approach even in late stages of COPD.

## Acknowledgements

The authors thank Christine Veith, Ingrid Breitenborn-Müller, Carmen Homberger, Miriam Wessendorf, Susanne Lich, Dagmar Fenner-Nau, Navya Sri Bonthu and Karin Quanz for their technical assistance. The authors thank Simone Kraut and Akylbek Sydykov for their support with echocardiography and Monika Heiner for assistance with FACS. Flow cytometry was performed by Gabriela Michel from the Flow Cytometry Core Facility of the Medical Faculty at Justus Liebig University Giessen using a BD FACSymphony S6 funded by DFG/EU.

## Author contributions

CGC, VBL, SH, OP, MPM and NS contributed to the study design, data analysis, and interpretation; CGC, VBL, SH, CN, DG, MH, LIG, BK, AS, SK, ML, SJ, MH, JK, MG, C-YW, LG, MH, CS, EL, AC, JB, UM, LP-O, JW, MB, BM, KT, LL, SH2, CS, SC and GK and NS were study investigators who collected and assessed the data; CGC, VBL, OP and NS drafted the manuscript; JW, MB, SH2, CS, GK, FG, HAG, WS, AG and NW critically reviewed the manuscript. All the authors reviewed and approved the final manuscript. CGC is designated as the first coauthor for performing all *in vivo* experiments, while VBL is designated as the second coauthor for her contributions to the *in vitro* experiments.

---

**Extended Data Figure 1.**
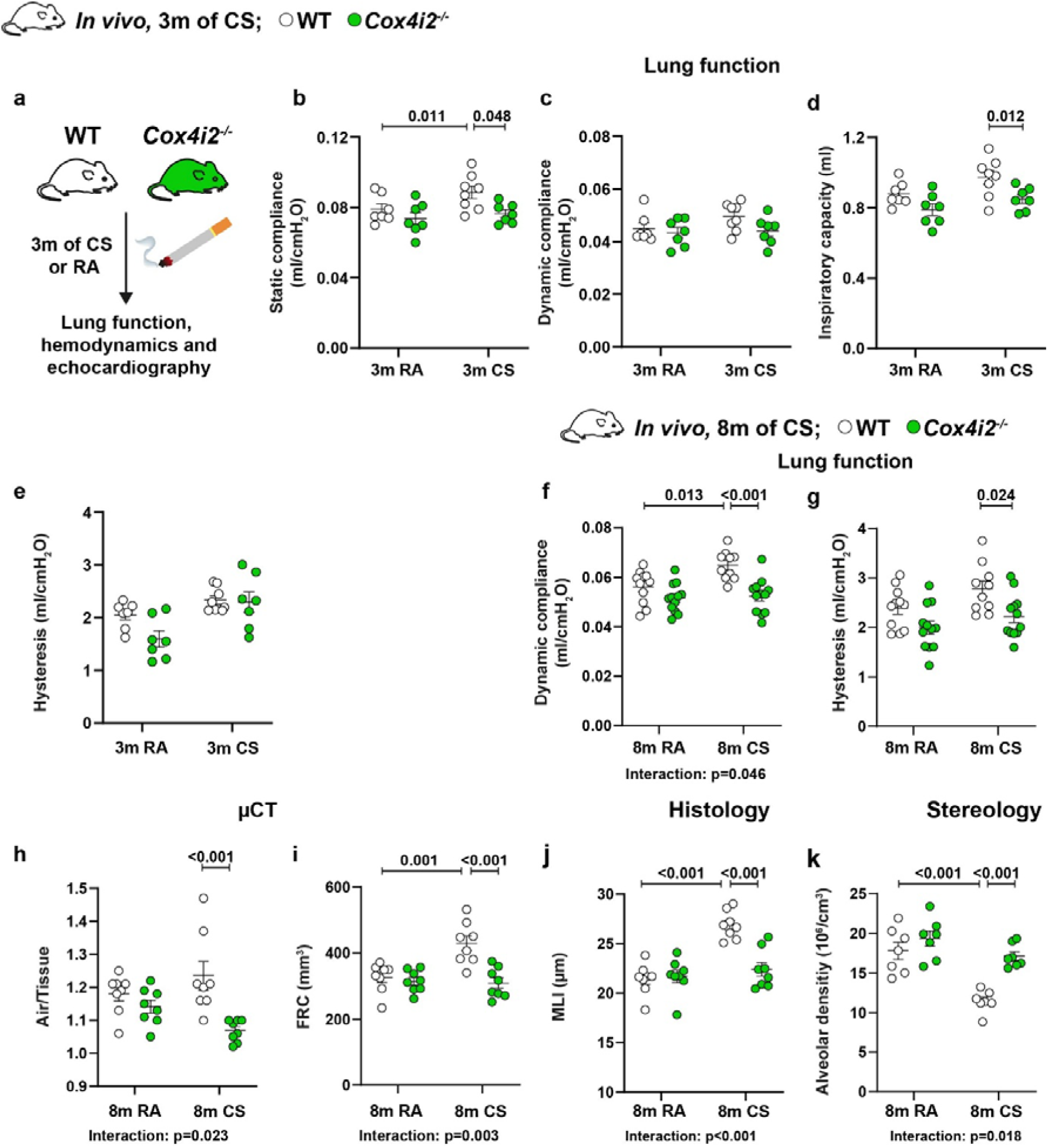
***Cox4i2* deficiency attenuates cigarette smoke-induced lung dysfunction** a) WT and *Cox4i2^-/-^* mice were exposed to either cigarette smoke (CS) or room air (RA) for 3 months. b-e) Lung function parameters after CS or RA exposure for 3 months (*n*=7–8 per group): static compliance (b), dynamic compliance (c), inspiratory capacity (d), and hysteresis (e). f, g) Lung function after CS or RA exposure for 8 months (n=10–12 per group): dynamic compliance (f) and hysteresis (g). h, i) µCT after CS or RA exposure for 8 months (n=8 per group): air/tissue ratio (h) and functional residual capacity (FRC) (i). j) Histological analysis of the lung parenchyma after CS or RA exposure for 8 months (n=8 per group): mean linear intercept (MLI). k) Stereological analysis of the lung parenchyma after CS or RA exposure for 8 months (n=7 per group): alveolar density. Statistical analysis was performed using two-way ANOVA. The data are presented as the mean ± SEM.

**Extended Data Figure 2.**
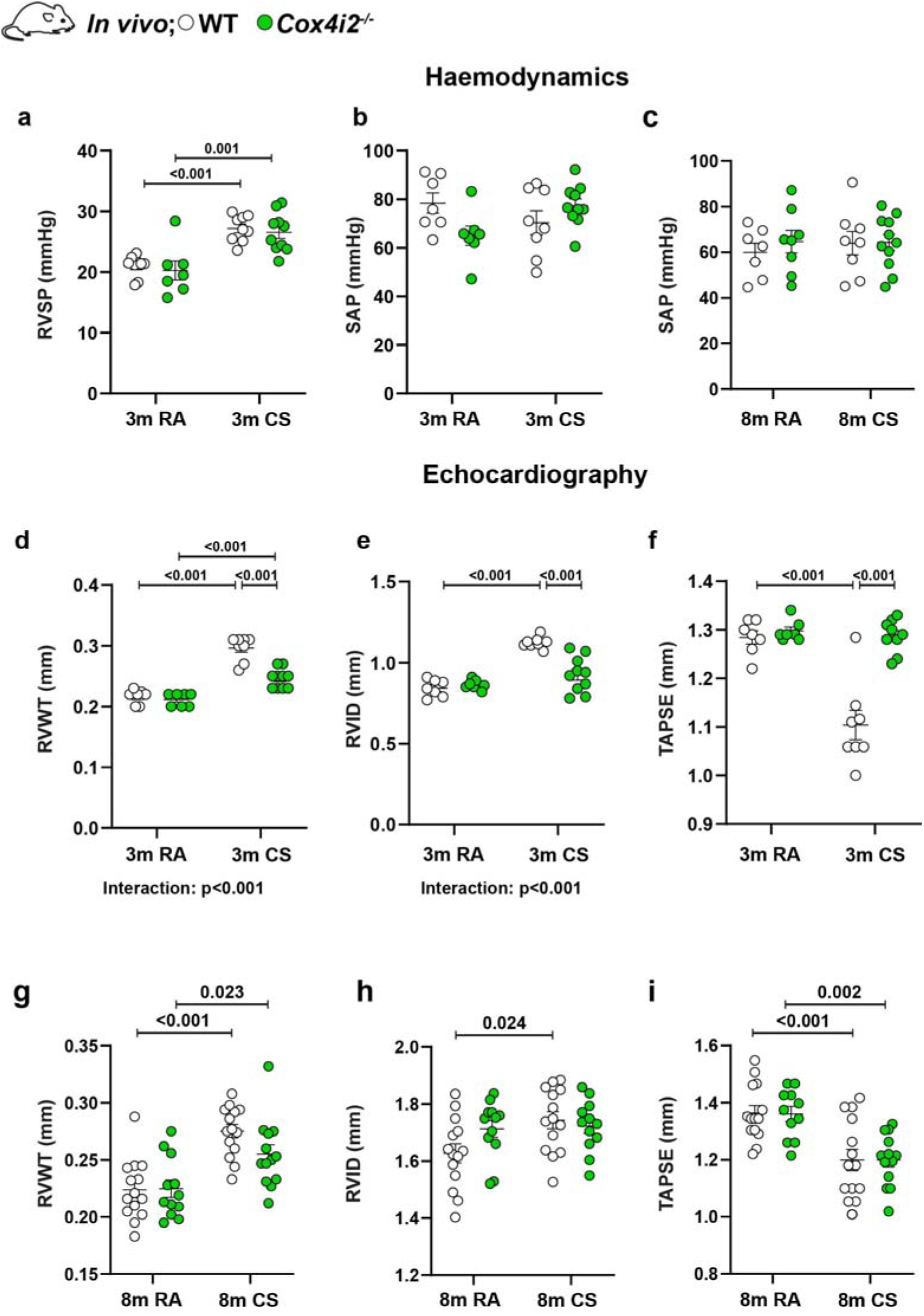
***Cox4i2* deficiency prevents the development of cigarette smoke-induced right ventricular remodelling but not pulmonary hypertension (PH).** a-c) Haemodynamic measurements after cigarette smoke (CS) or room air (RA) for 3 months of WT and *Cox4i2^-/-^* mice (*n*=7–10 per group): Right ventricular systolic pressure (RVSP) (a) and systolic arterial pressure (SAP) (b). c) SAP after CS or RA exposure for 8 months (*n*=7–11 per group). d-i) Echocardiographic analysis after CS or RA exposure for 3 months (d-f, *n*=7–10 per group) or 8 months (g-i, *n*=11-14 per group): right ventricular wall thickness (RVWT) (d, g), right ventricular internal diameter (RVID) (e, h) and tricuspid annular plane systolic excursion (TAPSE) (f, i). Statistical analysis was performed using two-way ANOVA. The data are presented as the mean ± SEM.

**Extended Data Figure 3.**
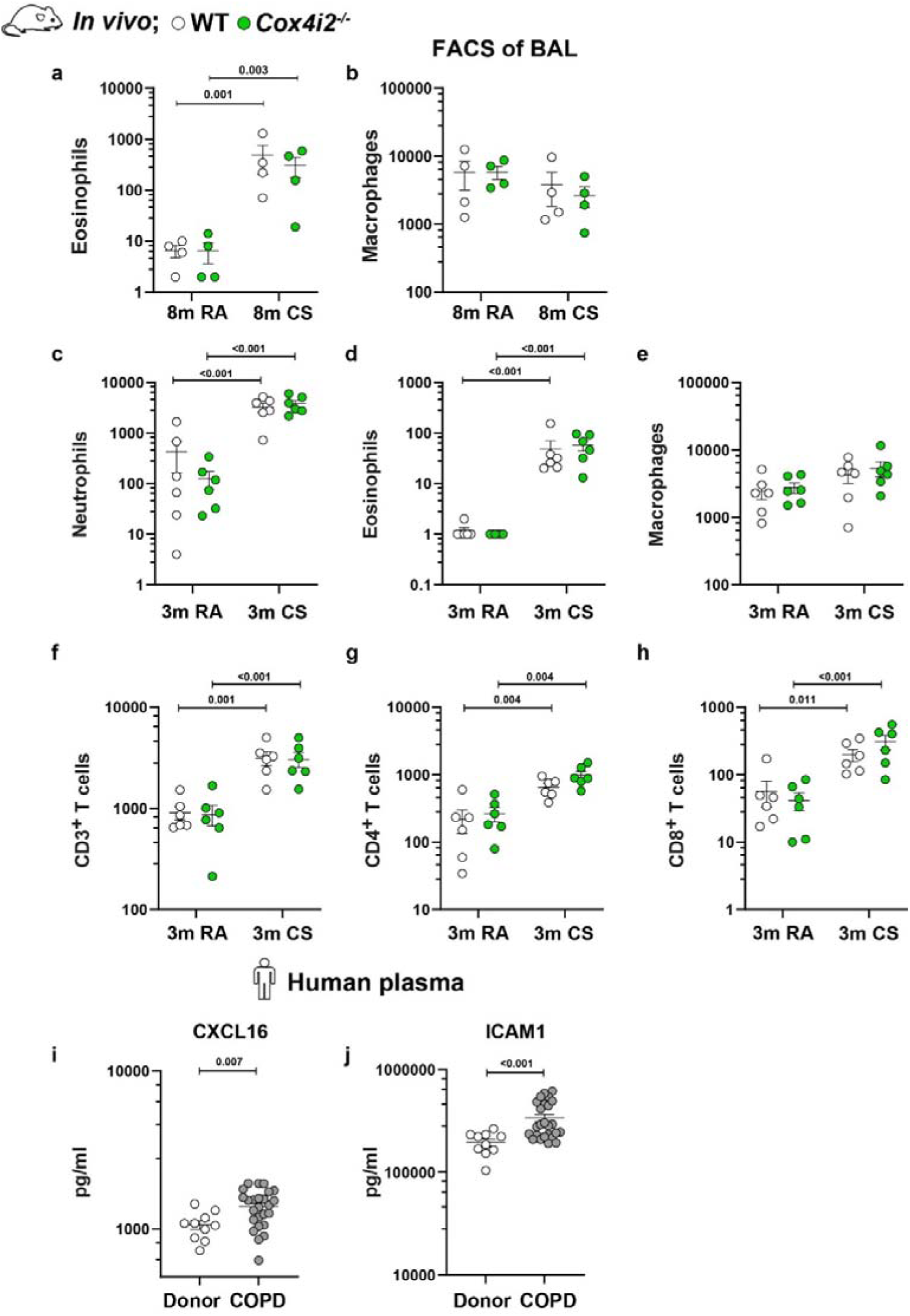
FACS analysis of bronchoalveolar lavage (BAL) and cytokines in human plasma. a, b) FACS analysis of bronchoalveolar lavage (BAL) from mice exposed to cigarette smoke (CS) or room air (RA) for 8 months (n=4 per group): eosinophils (a) and macrophages (b). c-h) FACS analysis of BAL from mice exposed to CS or RA for 3 months (m) (*n*=6 per group): Numbers of neutrophils (c), eosinophils (d), macrophages (e), CD3^+^ T cells (f), CD4^+^ T cells (g) and CD8^+^ T cells (h). i, j) The levels of CXCL16 (i) and ICAM1 (j) in plasma from donors (*n*=10) and patients with COPD (*n*=25). Statistical analysis was performed using an unpaired Student’s *t* test and two-way ANOVA. Data were log-transformed prior to statistical analysis. The data are presented as the mean ± SEM.

**Extended Data Figure 4.**
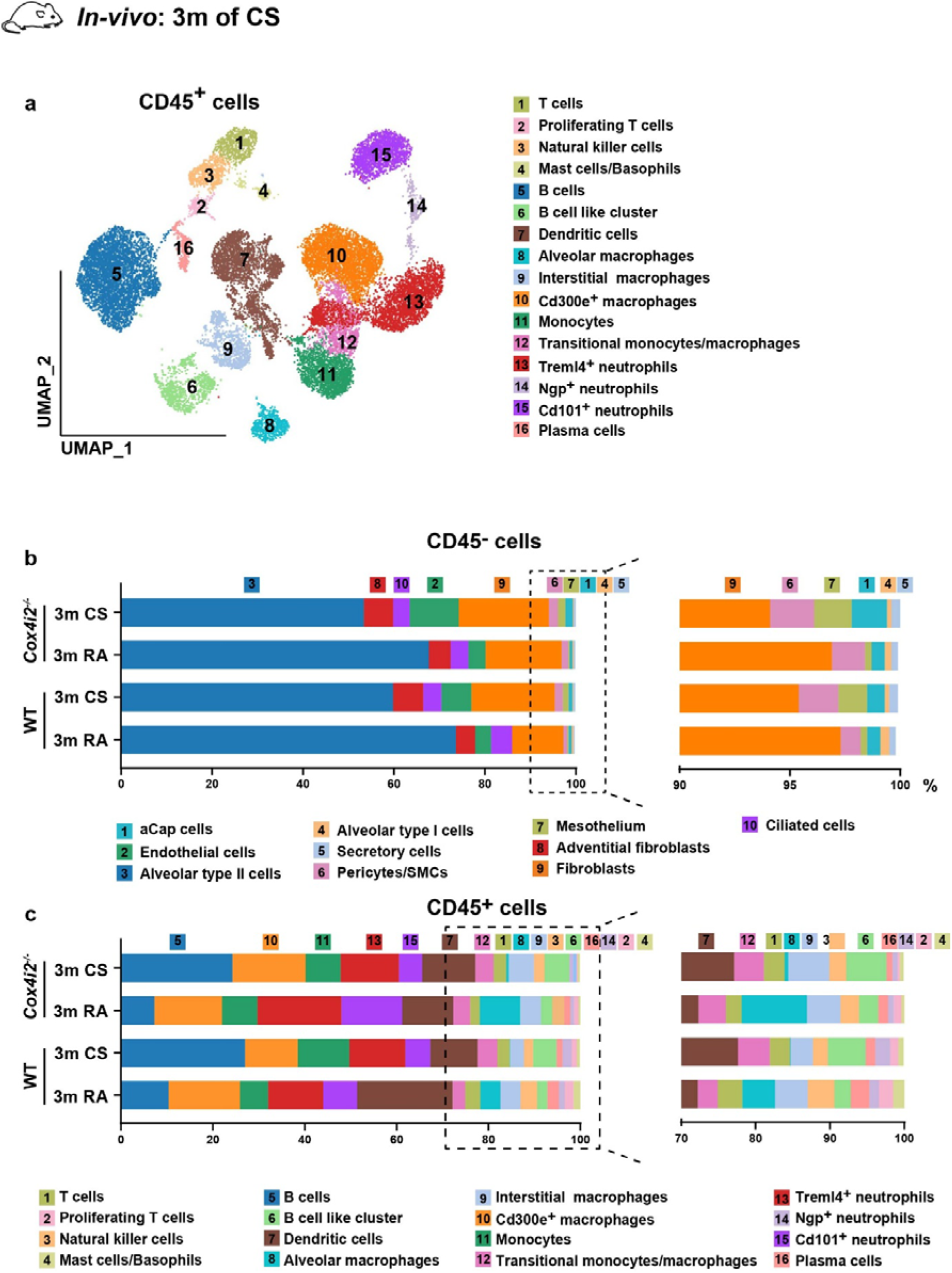
**Cell clusters identified by scRNA-seq of CD45^-^ and CD45^+^ lung cells after 3 months of CS exposure** a) UMAP representation of CD45^+^ cells (n=2–3 per group). b, c) Numbers of different clusters of CD45^-^ (b) and CD45^+^ (c) cells.

**Extended Data Figure 5.**
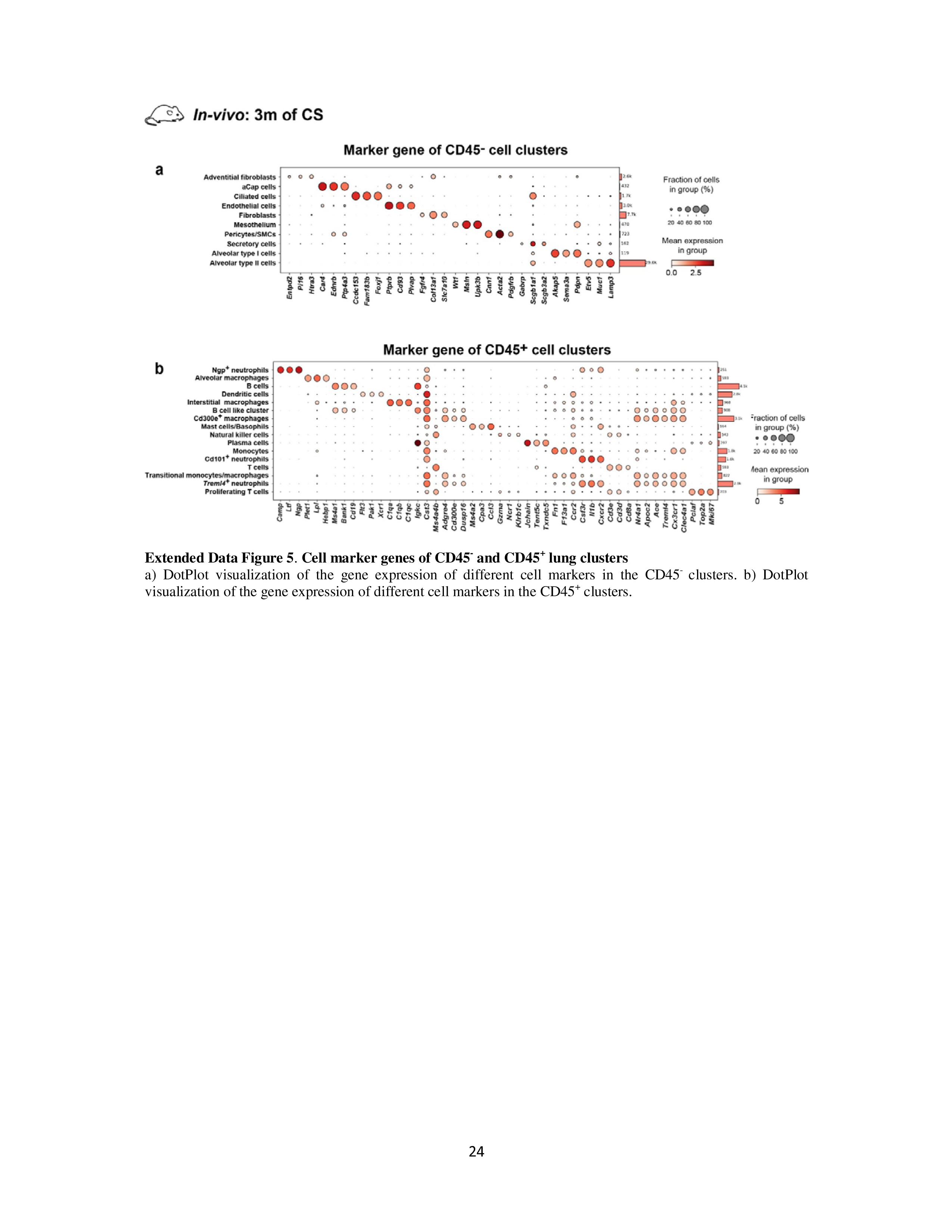
**Cell marker genes of CD45^-^ and CD45^+^ lung clusters** a) DotPlot visualization of the gene expression of different cell markers in the CD45^-^ clusters. b) DotPlot visualization of the gene expression of different cell markers in the CD45^+^ clusters.

**Extended Data Figure 6.**
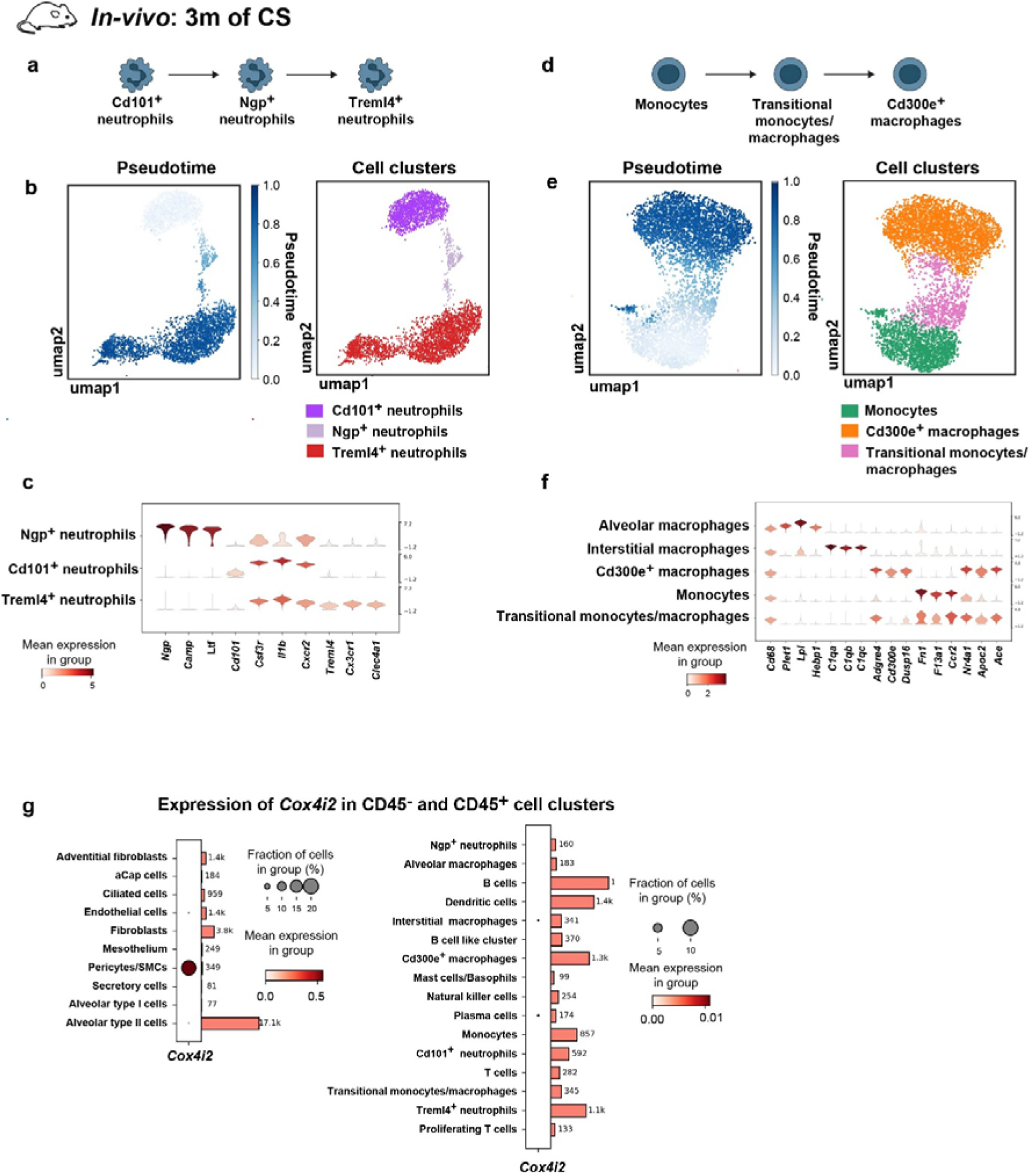
**Expression of *Cox4i2* in different cell clusters identified by scRNA-seq of CD45^-^and CD45^+^ lung cells** a-c) Annotation strategy of neutrophils at different developmental stages (a) by analysing pseudotime trajectory (b) and violin plots of gene expression of different neutrophil markers (c): Cd101^+^ neutrophils (*Cd101, Csf3r,* and *Il1b*), Ngp^+^ neutrophils (*Ngp, Ltf,* and *Camp*), and Treml4+ neutrophils (*Treml4, Cx3cr1,* and *Clec4a1*). d-f) Schematic illustration of the developmental stage of monocytes (d); pseudotime trajectory of monocytes (e) and violin plots of monocyte and macrophage markers (f). Monocyte/macrophage clusters expressing the classical marker *Cd68* were classified on the basis of distinct marker expression: alveolar macrophages (*Plet1, Lpl,* and *Hebp1*), interstitial macrophages (*C1qa, C1qb,* and *C1qc*), Cd300e^+^ macrophages (*Ace, Cd300e,* and *Dusp16*), monocytes (*Fn1, F13a1,* and *Ccr2*), and transitional monocyte/macrophage clusters (*Nr4a1, Apoc2,* and *Clec4a1*), which coexpress markers of nonclassical monocytes and maladaptive macrophages. g) Expression of *Cox4i2* mRNA across different cell clusters within CD45^-^ and CD45^+^ cell populations.

**Extended Data Figure 7.**
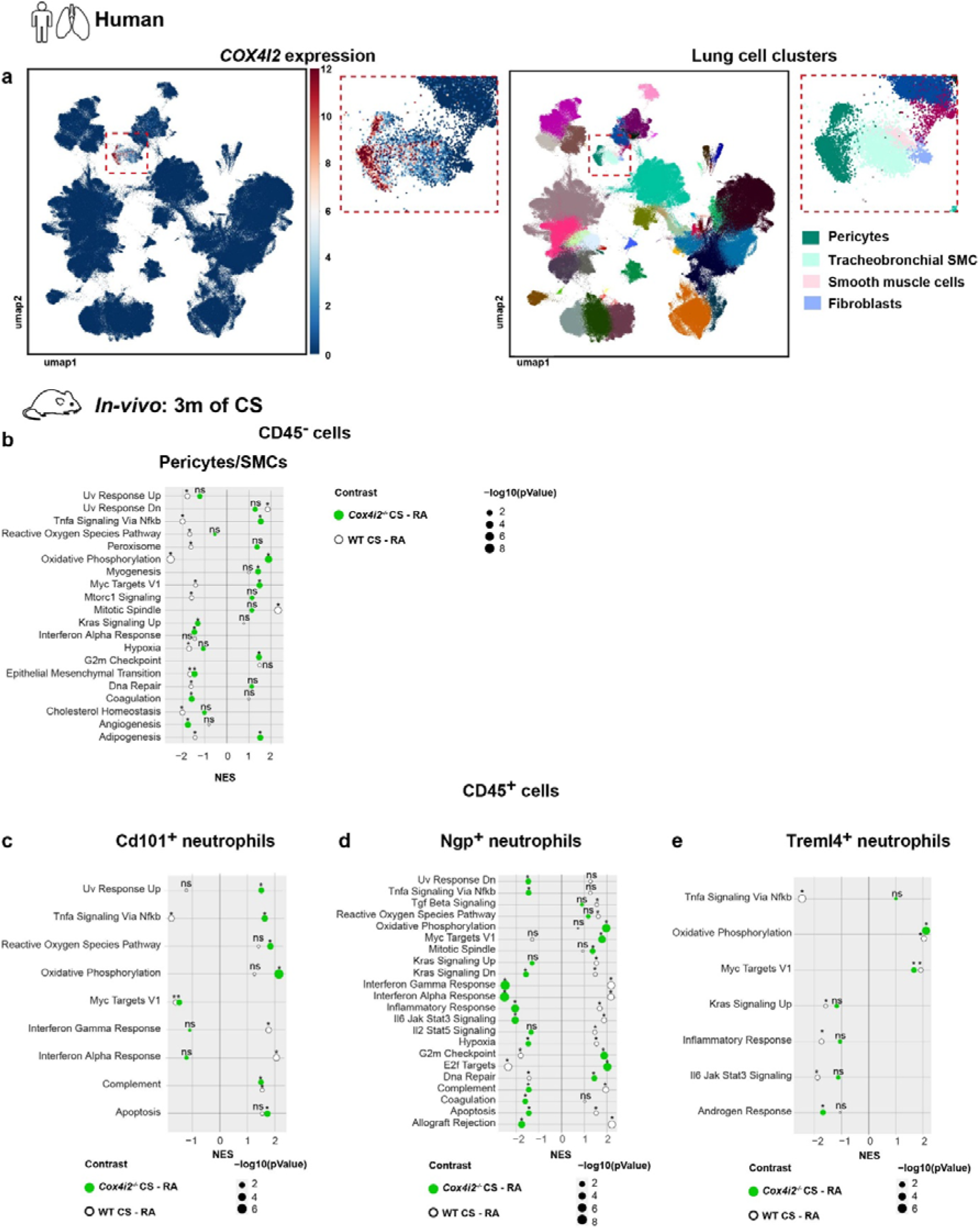
**Expression of *COX4I2* in different cell clusters of human lung cells and enrichment analysis of CD45^-^ and CD45^+^ mouse lung cell clusters.** a) UMAP representation of *COX4I2* expression in human lung cells. b) Hallmark gene set enrichment analysis of the Pericyte/SMC cluster in CD45^-^ cells. c-e) Hallmark gene set enrichment analysis of Cd101^+^ neutrophils (c), Ngp^+^ neutrophils (d) and Treml4^+^ neutrophils (e) of CD45^+^ cells.

**Extended Data Figure 8.**
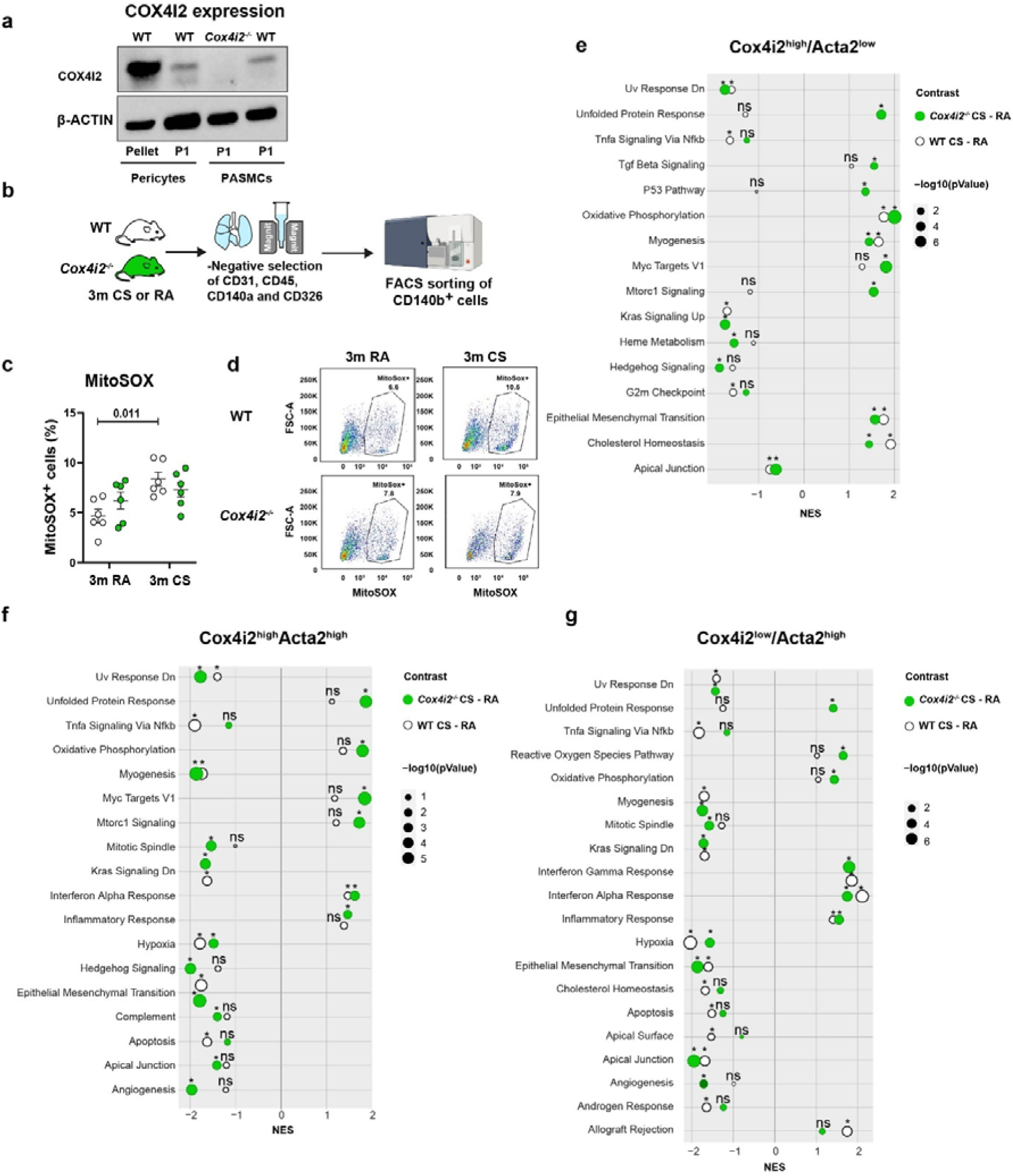
**COX4I2 expression in pericytes and ACTA2^+^ PCs, level of mitochondrial ROS in pericytes exposed to cigarette smoke for 3 months and enrichment analysis of lung pericytes single cell sequence data** a) Western blot analyses of COX4I2 protein expression in the pellet of pericytes directly after isolation, and in cultured pericytes and PASMCs. PASMCs isolated from *Cox4i2*^−/−^ mice used as negative control. b) Schematic procedure for isolating pericytes from *Cox4i2*^−/−^ and WT mice after 3 months of exposure to cigarette smoke (CS) or room air (RA). c, d) MtROS levels in pericytes from *Cox4i2*^−/−^ and WT mice after exposure to CS or RA for 3 months (*n*=6 individual cell isolations per group). c: Quantification, d: Representative FACS staining of pericytes with 5 µM MitoSOX. Statistical analysis was performed using two-way ANOVA. The data are presented as the mean ± SEM. e-g) Hallmark gene set enrichment analysis of subclusters of pericytes.

**Extended Data Figure 9.**
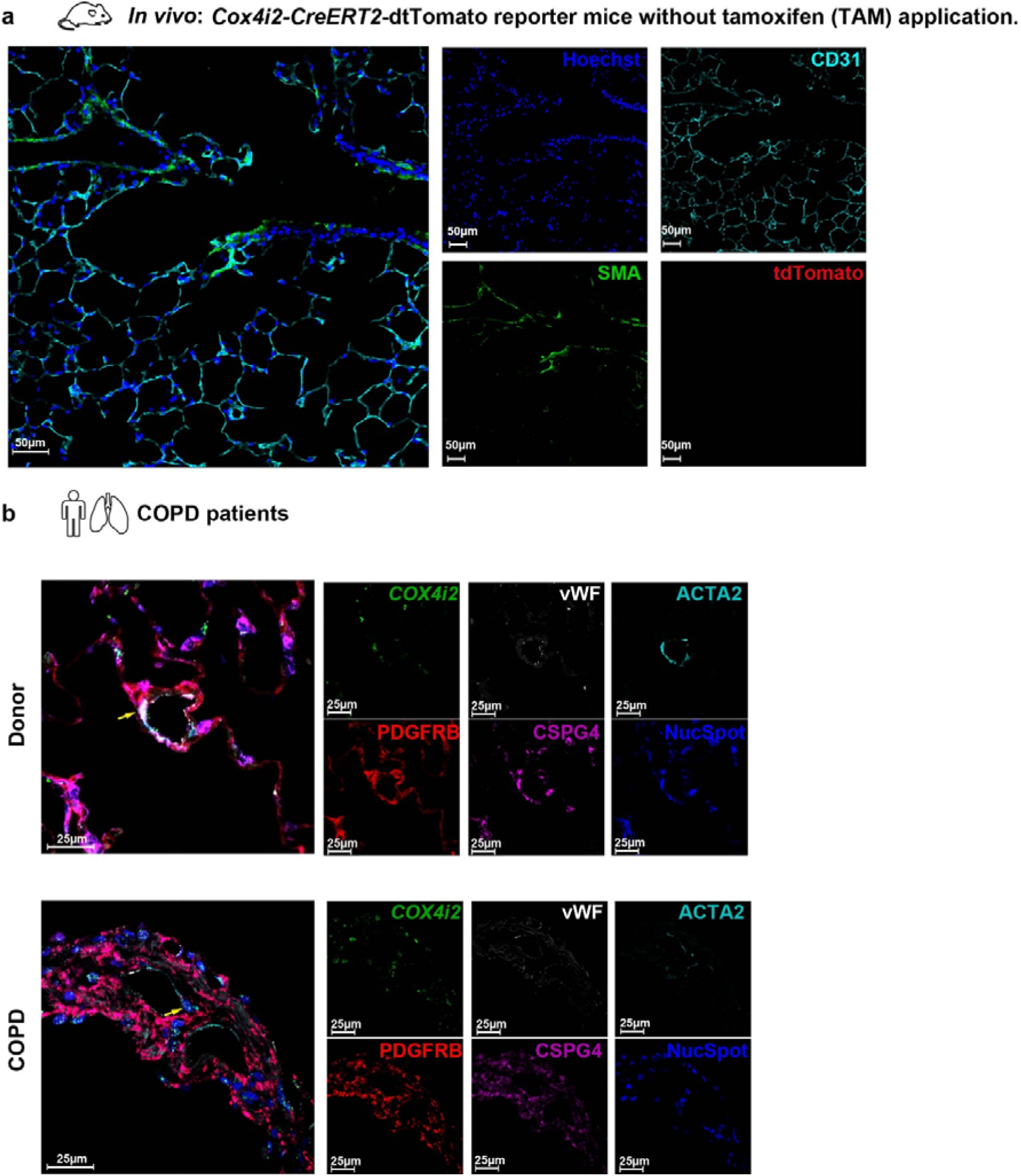
***COX4I2* mRNA expression in human lungs** a) Representative images of lung tissue from *Cox4i2*-CreERT2-tdTomato mice without tamoxifen (TAM) application. Colour code: red – tdTomato, green – smooth muscle actin, bright cyan – CD31, blue – Hoechst. b) *In situ* hybridization for the localization of *COX4I2* mRNA in healthy (donor) and COPD lungs. Coimmunofluorescence staining for pericyte (PDGFRB and CSPG4), smooth muscle (ACTA2), and endothelial (Von Willebrand factor - vWF) markers. Colour code: green - *COX4I2* mRNA, white – vWF, bright cyan – ACTA2, red – PDGFRB, magenta - CSPG4 and blue – NucSpot.

**Extended Data Figure 10.**
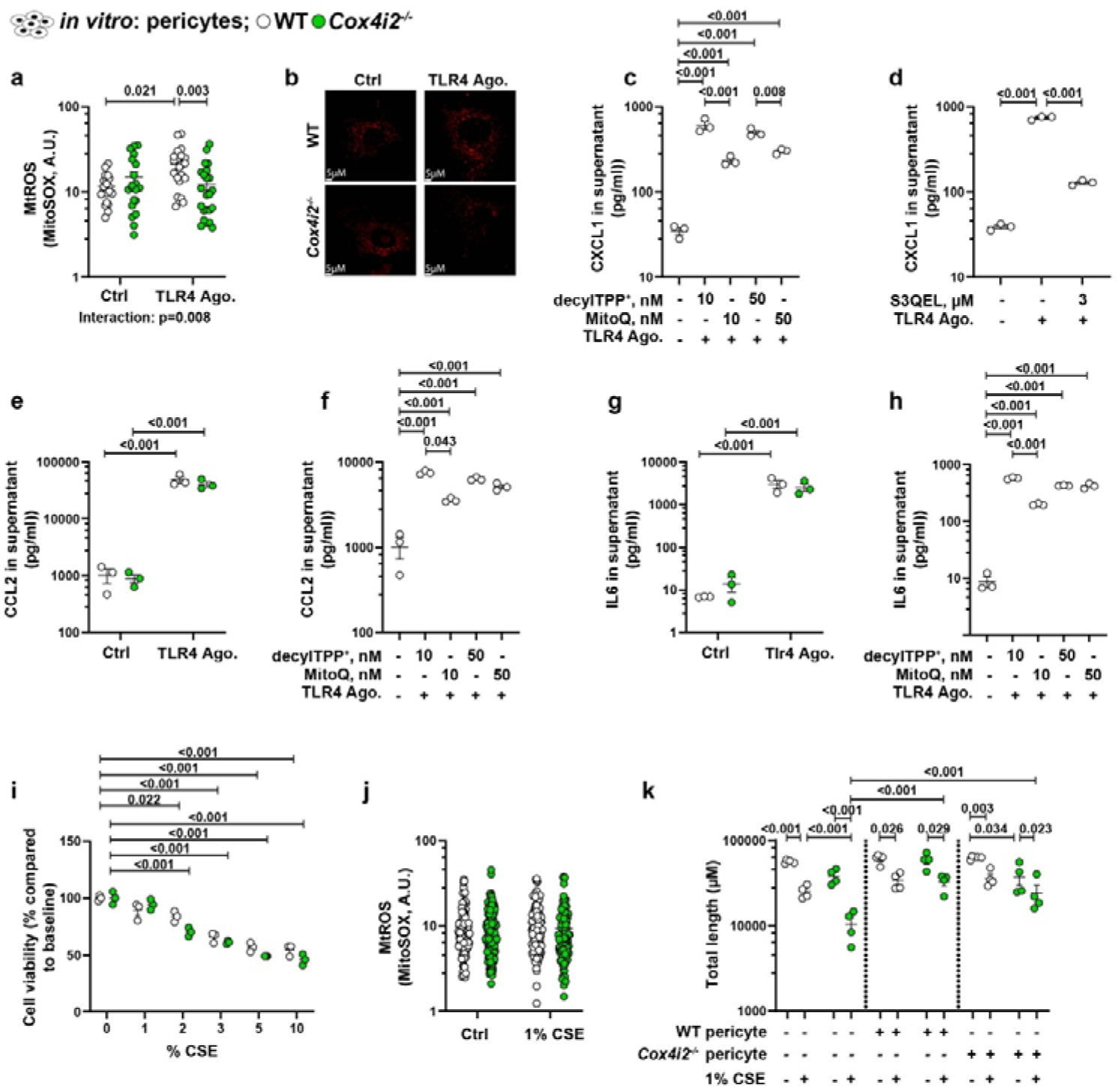
**Effects of S3QEL on CXCL1 levels, *Cox4i2* deficiency on chemokines/cytokine levels in cell culture supernatants from pericytes and mitochondrial ROS production in pericytes** a-b) MtROS production in pericytes after stimulation with the TLR4 agonist LPS-EB, as measured by MitoSOX. *n*=20–23 cells from three independent cell isolations per group. c, d) Protein expression of CXCL1 in cell culture supernatant from WT pericytes after stimulation with the TLR4 agonist LPS-EB following the application of MitoQ or its inactive carrier, DecylTPPLJ (c), or the mitochondrial complex III superoxide inhibitor 3 µM S3QEL (d). *n*=3 individual cell isolations per group. e, f) The protein level of CCL2 in the cell culture supernatant isolated from *Cox4i2*^−/−^ and WT mice (e) or from WT mice following treatment with MitoQ or its inactive carrier, DecylTPPLJ (f), and stimulation with the TLR4 agonist LPS-EB. *n*=3 individual cell isolations per group. g, h) Protein level of IL6 in the cell culture supernatant isolated from *Cox4i2*^−/−^ and WT mice (g) or from WT mice following treatment with MitoQ or its inactive carrier, DecylTPPLJ (h), and stimulation with the TLR4 agonist LPS-EB. *n*=3 individual cell isolations per group. i) Cell viability of primary pulmonary pericytes isolated from *Cox4i2*^−/−^ and WT mice and exposed to various doses of cigarette smoke extract (CSE). *n*=3 independent cell isolation per group. j) Mitochondrial ROS production in pericytes after exposure to 1% CSE measured by MitoSOX. *n*=104–132 cells from three independent cell isolations per group. k) Vascular tube formation of ECs incubated with solvent CSE in the presence or absence of primary pulmonary pericytes. *n*=4 isolation per group: tube length. Statistical analysis was performed using one-way or two-way ANOVA. Data from panels a, c-h and j were log-transformed prior to statistical analysis. The data are presented as the mean ± SEM.

**Extended Data Figure 11.**
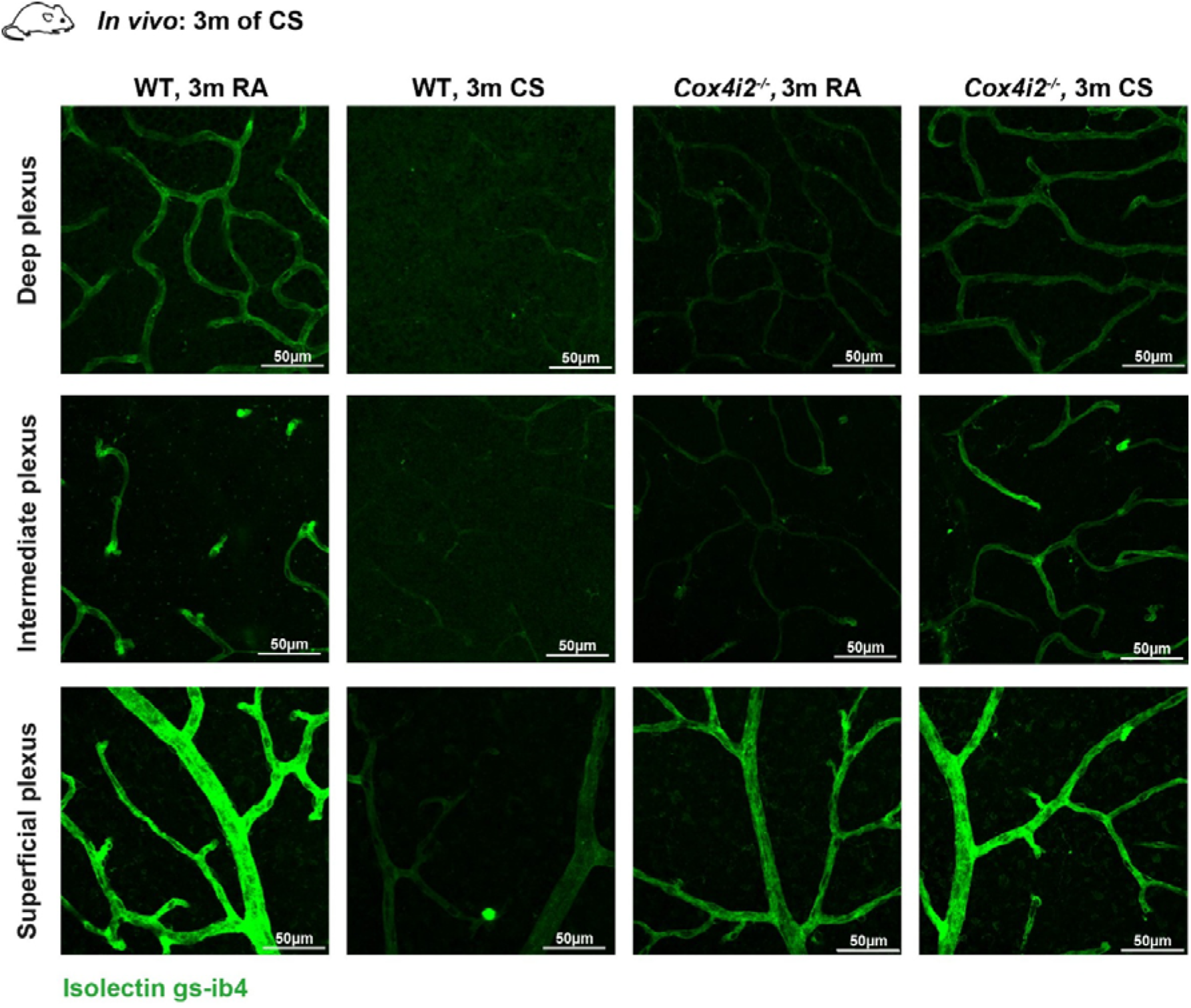
***Cox4i2* deficiency reduces capillary network density under basal conditions but prevents the detrimental effects of cigarette smoke** The capillary network of the superficial, intermediate, and deep plexuses was visualized by fluorescent labelling using isolectin B4 (green) in WT and *Cox4i2^-/-^* mice exposed to either cigarette smoke (CS) or room air (RA) for 3 months.

**Extended Data Figure 12.**
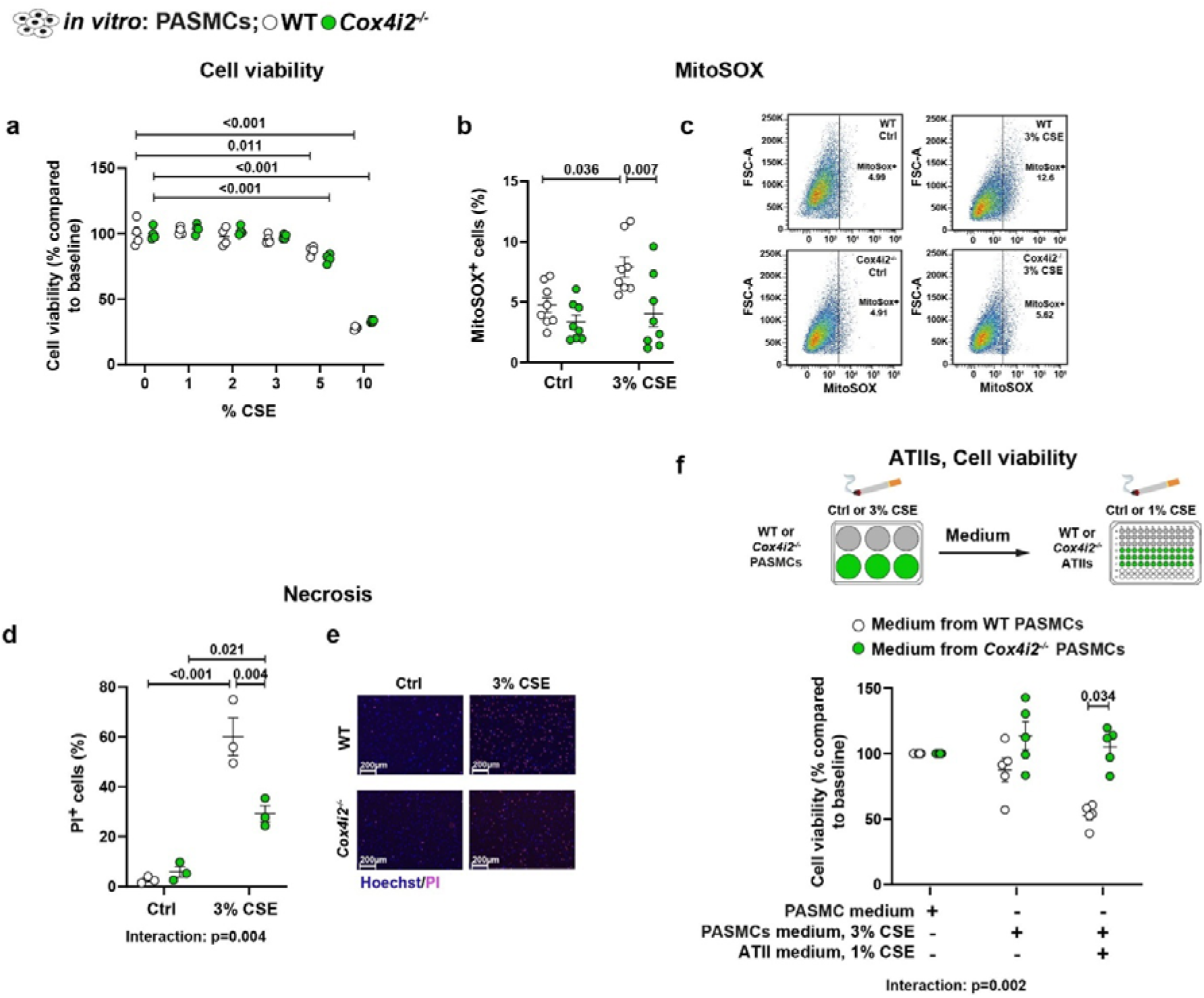
***In vitro* effects of *Cox4i2* deficiency in pericytes and PASMCs** a) Viability of PASMCs isolated from *Cox4i2^-/-^* and WT mice and exposed to various doses of cigarette smoke extract (CSE) (*n*=4 individual cell isolations per group). b, c) Mitochondrial ROS production in PASMCs after CSE exposure determined by MitoSOX (*n*=8 individual cell isolations per group). b: Quantification, c: representative FACS staining of PASMCs with 5 µM MitoSOX. d, e) Necrosis of PASMCs determined by propidium iodide (PI) staining (*n*=3 individual cell isolations per group), d: Quantification, e: Representative pictures. f) ATII cell viability after exposure to 1% CSE in the presence of medium from WT or *Cox4i2*^−/−^ PASMCs, which were exposed to solvent (PASMC medium) or 3% CSE (PASMC medium, 3% CSE). *n*=5 individual cell isolations per group. Statistical analysis was performed using two-way ANOVA. The data are presented as the mean ± SEM.

**Extended Data Figure 13.**
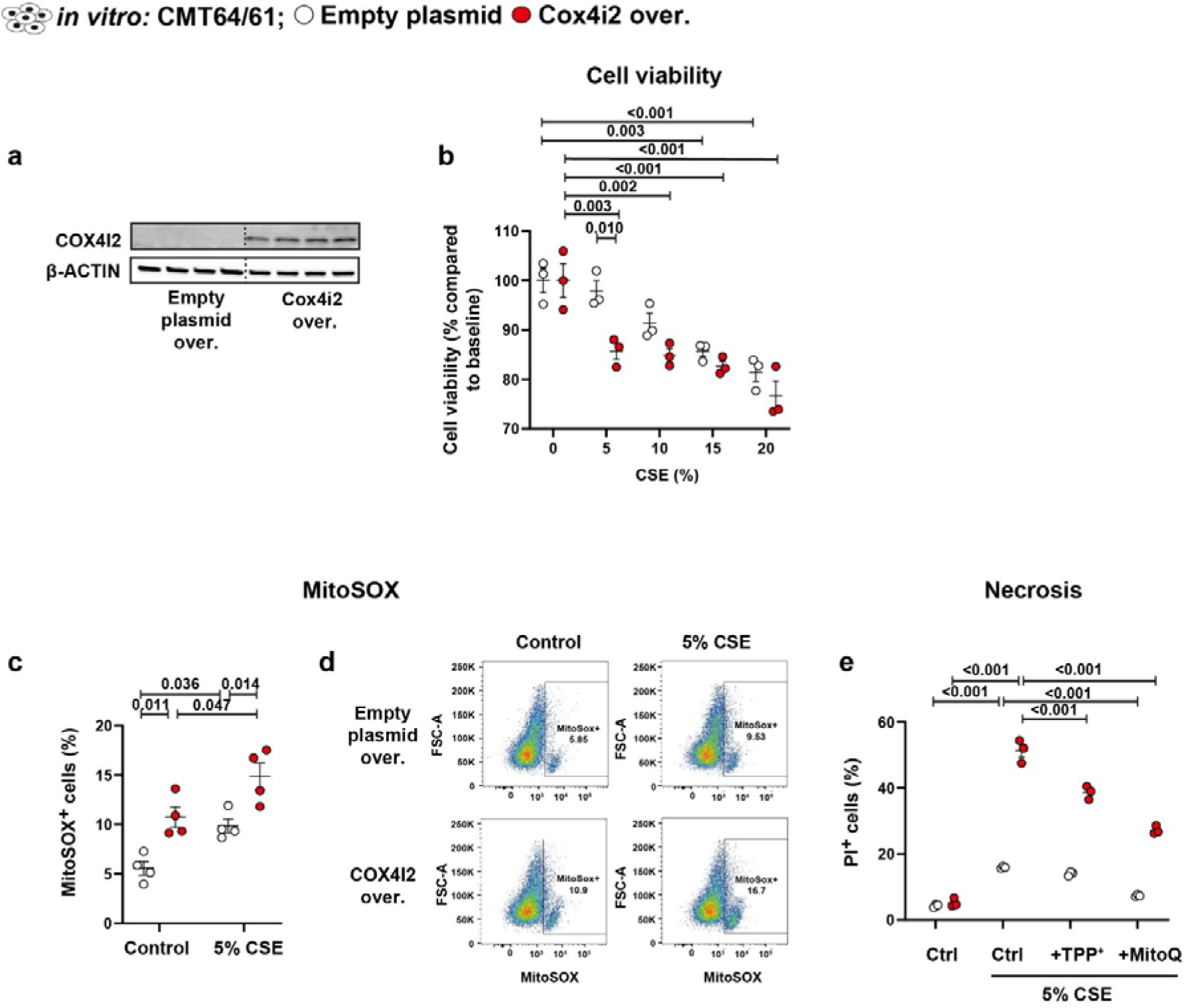
***In vitro* effects of *Cox4i2* overexpression in CMT64/61 cells** a) Western blot analysis of COX4I2 protein expression in CMT64/61 cells after expression of the empty plasmid or *Cox4i2*-expressing plasmid. b) Viability of CMT64/61 cells exposed to various doses of CSE (*n*=3 per group). c, d) Mitochondrial ROS production in CMT64/61 cells after CSE exposure determined by MitoSox (*n*=4 per group). c: Quantification, d: Representative FACS staining of CMT64/61 cells with 5 µM MitoSOX. e) Necrosis of CMT64/61 was determined by propidium iodide (PI) staining (*n*=3 per group), and statistical analysis was performed using two-way ANOVA. The data are presented as the mean ± SEM.

**Extended Data Figure 14.**
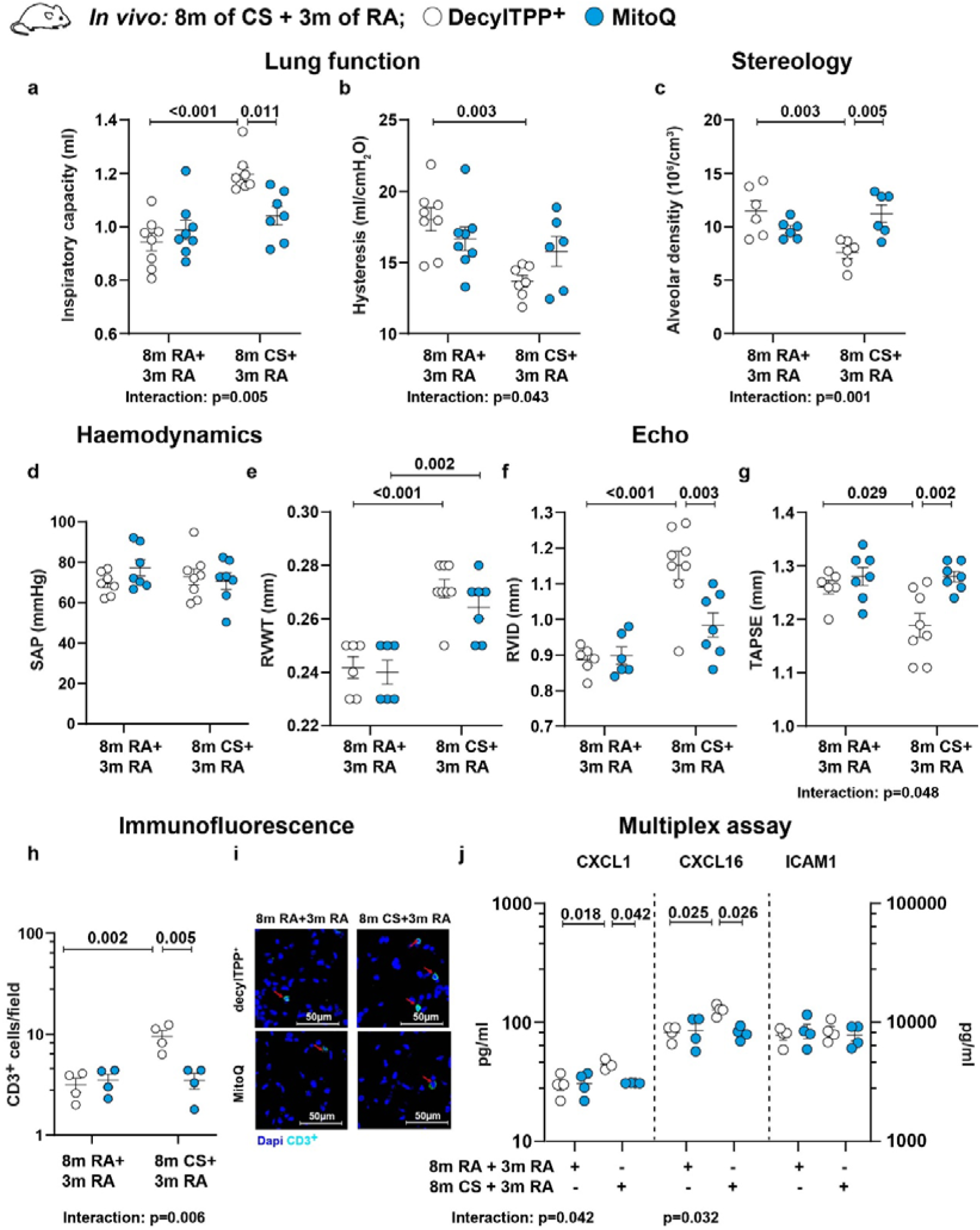
**MitoQ treatment reverses emphysema, right ventricular remodelling and dysfunction after 8 months of cigarette smoke exposure and cytokine analysis of bronchoalveolar lavage fluid (BALF) from WT mice exposed to cigarette smoke for 8 months and treated with MitoQ for 3 months** a, b) Lung functional analyses (*n*=6-8 per group) of inspiratory capacity (a) and hysteresis (b). c) Stereological analysis of the lung parenchyma (n=6 per group): alveolar density. d) Haemodynamic measurements (n=7–8 per group): systolic arterial pressure (SAP) e-g) Echocardiographic analysis (*n*=6–8 per group) of right ventricular wall thickness (RVWT) (e), right ventricular internal diameter (RVID) (f) and tricuspid annular plane systolic excursion (TAPSE) (g). h, i) Immunofluorescence analysis of CD3^+^ cell accumulation in the lung parenchyma (*n*=4 per group). h: Quantification. i: Representative images. g) Multiplex analysis of CXCL1, CXCL16 and ICAM1 expression in the BALF (*n*=4 per group). Statistical analysis was performed using two-way ANOVA. Data from panel h, g were log-transformed prior to statistical analysis. The data are presented as the mean ± SEM.

**Extended Data Table 1.**
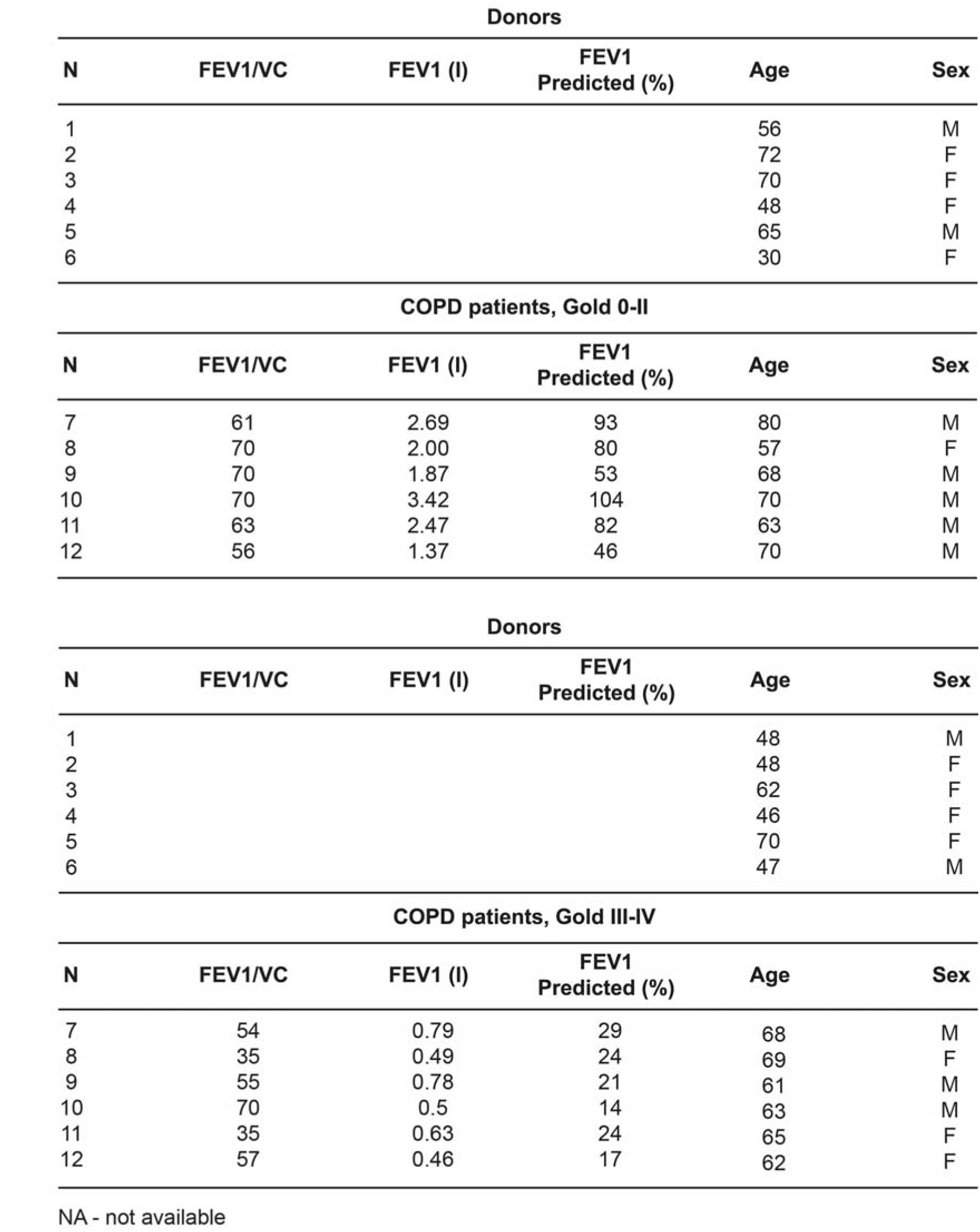
Patient characteristics of samples from donor/COPD lung.

| Donors |  |  |  |  |  |
| --- | --- | --- | --- | --- | --- |
| N | FEV1/VC | FEV1 (l) | FEV1 Predicted (%) | Age | Sex |
| 1 |  |  |  | 56 | M |
| 2 |  |  |  | 72 | F |
| 3 |  |  |  | 70 | F |
| 4 |  |  |  | 48 | F |
| 5 |  |  |  | 65 | M |
| 6 |  |  |  | 30 | F |
| COPD patients, Gold 0-II |  |  |  |  |  |
| N | FEV1/VC | FEV1 (l) | FEV1 Predicted (%) | Age | Sex |
| 7 | 61 | 2.69 | 93 | 80 | M |
| 8 | 70 | 2.00 | 80 | 57 | F |
| 9 | 70 | 1.87 | 53 | 68 | M |
| 10 | 70 | 3.42 | 104 | 70 | M |
| 11 | 63 | 2.47 | 82 | 63 | M |
| 12 | 56 | 1.37 | 46 | 70 | M |
| Donors |  |  |  |  |  |
| N | FEV1/VC | FEV1 (l) | FEV1 Predicted (%) | Age | Sex |
| 1 |  |  |  | 48 | M |
| 2 |  |  |  | 48 | F |
| 3 |  |  |  | 62 | F |
| 4 |  |  |  | 46 | F |
| 5 |  |  |  | 70 | F |
| 6 |  |  |  | 47 | M |
| COPD patients, Gold III-IV |  |  |  |  |  |
| N | FEV1/VC | FEV1 (l) | FEV1 Predicted (%) | Age | Sex |
| 7 | 54 | 0.79 | 29 | 68 | M |
| 8 | 35 | 0.49 | 24 | 69 | F |
| 9 | 55 | 0.78 | 21 | 61 | M |
| 10 | 70 | 0.5 | 14 | 63 | M |
| 11 | 35 | 0.63 | 24 | 65 | F |
| 12 | 57 | 0.46 | 17 | 62 | F |

NA - not available

**Extended Data Table 2.**
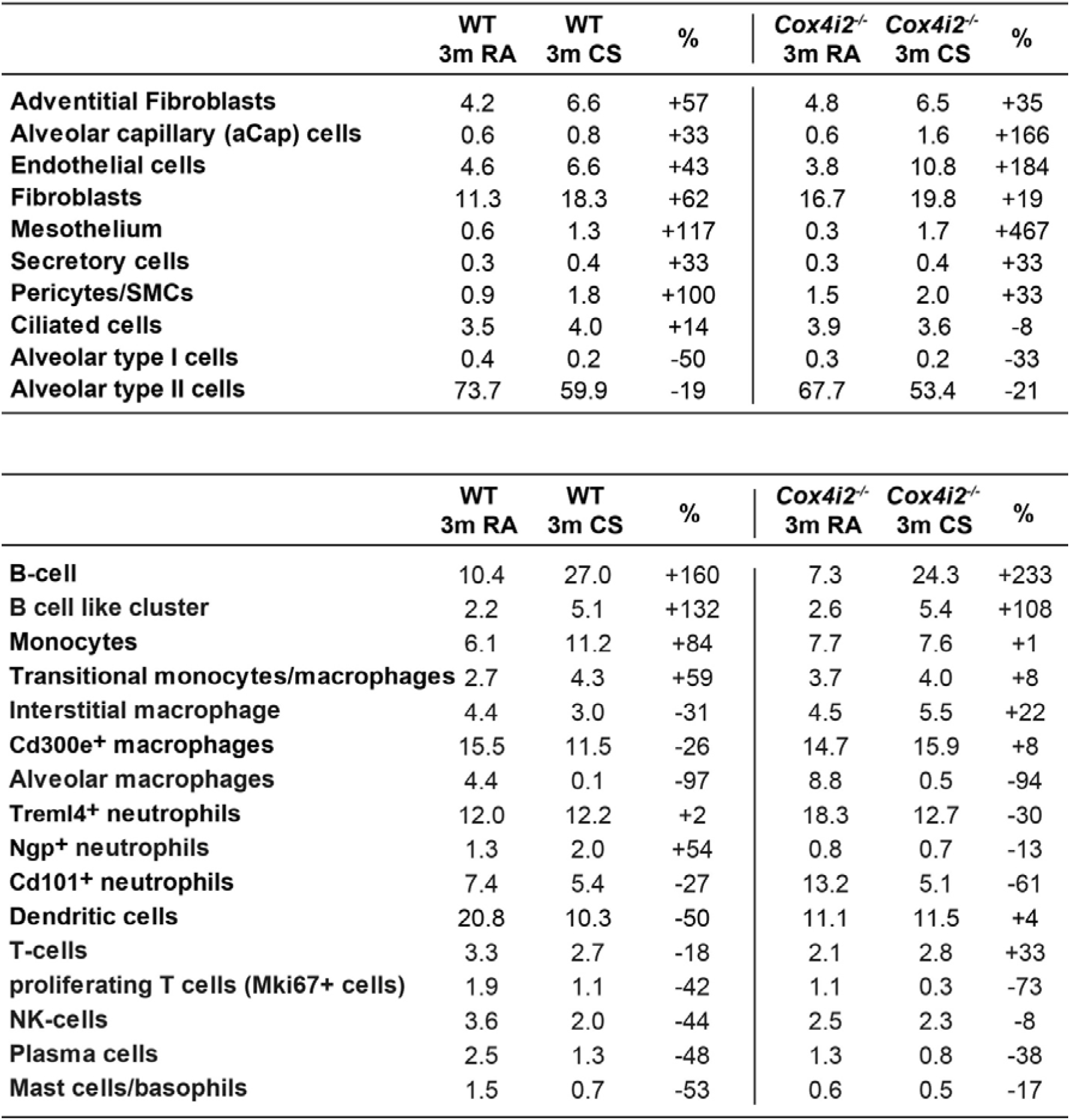
Percentage of cells in CD45+ and CD- cell clusters.

## Extended Data Video 1

Labelling of *Cox4i2*-expressing cells was performed using *Cox4i2*-CreERT2-tdTomato mice. A Kozak-CreERT2-P2A cassette was inserted just before the *Cox4i2* start codon, and these mice were crossed with tdTomato reporter mice containing a loxP-STOP-loxP cassette. After tamoxifen (i.p.; TAM) injection, Cre activation removed the stop cassette, enabling tdTomato expression driven by the *Cox4i2* promoter. Colour code: Red – tdTomato, green – SMC actin, white – CD31, blue – Hoechst.

## Methods

### Animals and Experimental Design

Animal experiments were conducted with strict adherence to the guidelines of Directive 2010/63/EU and were approved by local authorities (Regierungspräsidium, Giessen, animal proposal number G17/2016, G70/2019, G84/2020, G56/2023 and G63/2024). *Cox4i2^-/-^*(B6.129 Cox4i2tm1Hut) mice, provided by Prof. Mike Hütteman, were bred in-house, while C57BL/6J (WT) mice served as controls^14^. Three-month-old male and female WT and *Cox4i2^-/-^* mice were randomly assigned to CS or RA exposure for 3 (n=10 per group) or 8 (n=14 per group) months. CS exposure (3R4F cigarettes, 140 mg particulate matter/m³, Kentucky Tobacco Research & Development Centre) occurred for 6 hours/day, 5 days/week. All mice underwent *in vivo* assessments (haemodynamics, echocardiography, lung function, µCT). Occasional technical issues (e.g., catheter dislocation) led to variations in final sample sizes. After exposure to CS or RA for 3 months and *in vivo* functional studies, WT and *Cox4i2^-/-^* mice were subsequently randomly assigned to FACS analysis of BAL cells (*n*=6 per group) or to immunofluorescence quantification of CD31^+^ cells (*n* = 4 per group). After exposure to CS or RA for 8 months, WT and *Cox4i2^-/-^*mice were subsequently randomly assigned to either *in vivo* FMT–µCT imaging using Annexin Vivo 760 (*n* = 4 per group) or *post mortem* analyses. *Post mortem* assessments included FACS analysis of BAL cells (*n* = 4 per group), multiplex chemokine and cytokine profiling of BALF (*n* = 5 per group), histological analysis (*n* = 8 per group), stereological analysis (*n* = 7 per group), morphological analysis of pulmonary vessels (*n* = 4 per group), quantification of 3-nitrotyrosine staining (n = 4 per group), and immunofluorescence quantification of CD45^+^ and CD3^+^ cells (*n* = 4 per group).

For vessel casting (*n*□=□3 per group), and single□cell sequencing of either total lung cells (CD45□ and CD45□, n□=□3 per group) or FACS-sorted pericytes (*n*□=□3 per group), WT and *Cox4i2^-/-^*mice were exposed to CS or RA for three months and subsequently subjected to the experiment without *in vivo* functional studies.

For the curative approach, MitoQ (mitoquinone mesylate, Antipodean Pharmaceuticals, New Zealand) treatment (50 mg/kg/day, intragastric by gavage) was initiated after 8 months of CS exposure and continued for 3 months, with CS discontinued. Controls included age-matched RA-exposed and placebo-treated (1-decyl)triphenylphosphonium bromide (decylTPP^+^). Only male mice were used for this study (n=10 per group).

After MitoQ or DecylTPP^+^ treatment, all mice underwent *in vivo* assessments and were subsequently randomly assigned to multiplex chemokine and cytokine profiling of BALF (*n* = 4 per group), stereological analysis (*n* = 6 per group), morphological analysis of pulmonary vessels (*n* = 5 per group), quantification of 3-nitrotyrosine staining (*n* = 4 per group), immunofluorescence quantification CD45^+^ and CD3^+^ cells (*n* = 4 per group).

To generate a mouse line expressing tdTomato under the control of the Cox4i2 promoter (Cox4i2-CreERT2-dtTomato), we inserted a Kozak-CreERT2-P2A cassette upstream of the ATG start codon located in exon 2 of the *Cox4i2* gene (Transcript: 201-ENSMUST00000010020). Genotyping was performed using a forward primer (5’-TTACCTCTAAGGCCTTTGCATCC-3’) and a reverse primer (5’-GTTTGTTGACACTAGAAGCAGGAGA-3’). The expected product size was 389 bp for the WT allele and 454 bp for the knock-in allele. These mice were then crossbred with B6;129S6-Gt(ROSA)26Sortm9(CAG-tdTomato)Hze/J mice (#:007905, the Jackson Laboratory), which contain a loxP-flanked STOP cassette preventing transcription of a CAG promoter-driven red fluorescent protein variant (tdTomato). All mice used in the experiments were heterozygous.

### Human samples

Human lung tissue samples were obtained from both healthy donors and patients with chronic obstructive pulmonary disease (COPD), and the studies were approved by the Ethics Committee of the Justus Liebig University School of Medicine (Approval Numbers: AZ 31/93, 10/06, 220/18).

### Cigarette smoke application in isolated perfused and ventilated mouse lungs

CS application in isolated perfused and ventilated mouse lungs was performed as described, previously^48^. Shortly, mice were deeply anesthetized with ketamine (100 mg/kg, i.p.) and xylazine (20 mg/kg, i.p.) following anticoagulation with heparin (2,500 U/kg, i.p.). Intubation was performed via a tracheostomy, and mice were ventilated with a normoxic gas mixture (21% O□, 5.3% CO□, balance N□, normobaric) using a piston pump (Minivent Type 845, Hugo Sachs Elektronik) at a tidal volume of 10 μl/g, 90 breaths/min, and a positive end-expiratory pressure of 3 cm H□O. The excised lung-heart complex was suspended to measure weight changes and perfused via the pulmonary artery with Krebs-Henseleit buffer using a peristaltic pump (ISM834A V2.10, Ismatec), while outflow perfusate was collected from the left atrium. CS was generated by burning a cigarette within one minute using a 1 L/min normoxic gas flow. CS was introduced into the lungs in deep breaths (3–4 seconds each) for 5 minutes, manually controlling the inspiratory pressure to prevent damage. This was repeated three times with one-hour intervals.

### Echocardiography, *in vivo* micro-Computed tomography (**_μ_**CT) and fluorescence molecular tomography (FMT micro-CT)

Transthoracic echocardiography was performed using the VEVO2100 system (VisualSonics) following established techniques^9,14,15^. μCT imaging utilized a Quantum GX µCT scanner (PerkinElmer) according to standard protocols^49^. FMT imaging was conducted with an FMT 2500 system (VisEn Medical) as previously described^9^.

### Lung function, haemodynamic assessment, alveolar and vascular morphometry, and stereological analysis

Lung function and haemodynamic assessments were conducted according to methods outlined in references^9,14,15,48^. Alveolar and vascular morphometry, along with stereological analyses, were performed using established procedures^9,48^.

### BAL collection and cell differential counts by FACS

Following euthanasia, bronchoalveolar lavage (BAL) was collected for analysis. The fluid was centrifuged, and the supernatant stored for cytokine analysis. Cells from the BAL were treated, counted, and prepared for FACS analysis. Flow cytometry was conducted using a BD LSR Fortessa, and data were analyzed with FlowJo software. The antibodies that were used are from Biolegend: CD45 (#103140), Ly6G (#127614), SiglecF (#552126), CD11c (#117306), CD11b (#Cd11b), CD40 (#124618), CD3 (#100236), CD4 (#100236), CD8 (#126616).

### Vessel casting

After death, mouse lungs were perfused through the pulmonary artery with a vasodilation buffer containing PBS, 4 mg/L papaverine, 1 g/L adenosine, and 1000 IU/L heparin, using a roller pump at 5 mL/min. Body temperature was kept at 37°C, with ventilation at 140–150 breaths/min. After exsanguination, the lungs were perfused with a 0.7% agarose and barium mixture at 0.5 mL/min, prepared and kept at 37°C. Lungs were then fixed by tracheal perfusion with formalin, embedded in 3% agarose, and imaged using a Quantum GX μCT scanner (PerkinElmer Inc.).

### Retinal blood vessels

Confocal images were captured using an Olympus FV10i confocal microscope, equipped with Argon and HeNe lasers. A high-resolution scanning of image stacks was performed with an UPlanSApo_60/1.35 (Olympus). For an analysis of the immunolabelled blood vessel plexus, a stack of 2–10 sections was used (1.0-µm z-axis step size). Blood vessel plexuses were reconstructed by collapsing the stacks into a single plane. Brightness and contrast of the final images were adjusted using Adobe Photoshop CS5 and all four groups investigated were processed with the same scale.

### Transcriptome Analysis by Microarray

Total RNA was extracted from lung tissue using the RNeasy Mini Kit (Qiagen). Amplified RNA was labelled with cyanine and hybridized onto 8×60K microarray slides (Agilent Technologies) for 22 hours at 65 °C. The slides were scanned with the InnoScan is900 at 2 μm resolution, and image analysis was performed using Mapix software. Data were processed using R software and the limma package, including log2 signal calculation, background correction, quantile normalization, and averaging of replicate spot signals, and were deposited in NCBI’s Gene Expression Omnibus (GEO) under GEO Series accession number and GSE314829. For genes with multiple probes, the probe with the highest average signal was chosen for analysis.

### Single-cell capture and library preparation

For scRNA-seq of whole lung cells, three WT and three *Cox4i2^-/-^*mice per group were exposed to CS for 3 months. After digestion, whole lung cells were sorted using MACS with biotin-conjugated antibodies against CD45 (Miltenyi Biotec) to isolate the CD45^+^ and CD45^-^fractions. For scRNA-seq of the pericyte-enriched fraction, three WT and three *Cox4i2^-/-^* mice per group were pooled after 3 months of CS exposure. Then, 8,000 cells were loaded into the Chromium Controller (10x Genomics, USA). The cDNA libraries were prepared according to manufacturer’s instructions. Following the 10x Genomics library preparation protocol, all samples were sequenced using an Illumina NovaSeq6000 sequencer. Demultiplexing and FASTQ file generation were carried out using Illumina’s bcl2fastq (version 2.19.0.316). Sequencing reads were mapped to the mouse mm10 reference genome (refdata-gex-mm10-2020-A, obtained from 10x Genomics) using STAR solo (version 2.7.9a), generating a UMI count matrix where cells (barcodes) correspond to columns and genes to rows. UMI deduplication was also performed during alignment with STAR. The STAR run was conducted in Velocyto mode to enable downstream velocity analysis. For droplet filtering, the EmptyDrops_CR parameter was applied.

### Analyses of single-cell RNA-seq data

Aligned reads were subjected to ambient RNA removal with CellBender (0.3.0)^50^. Resulting UMI count matrices were further pre-processed for downstream analysis using Scanpy (version 1.11.1) in a Python (3.10.17) environment^51^. Doublets were identified using Scrublet with default parameters^52^. To remove low-quality cells, doublets and cell debris specific filtering criteria were applied: cells were required to have between 1,500 and 40,000 UMI counts, with mitochondrial reads comprising no more than 15% of total counts. Additionally, each cell had to express at least 1,000 genes (with at least one mapped read per gene). To streamline the gene set, only genes expressed in a minimum of 20 cells were retained.

UMI counts per cell were normalized in respect to total counts over all genes, resulting in the same number of UMIs in every cell. Thereupon counts were log transformed. One mouse *Cox4i2^-/-^*CS in CD45^-^ group was excluded from the analysis due to having too few cells. Principal component analysis (PCA) was conducted for dimensionality reduction and data integration using Harmony^53^. A k-nearest-neighbors approach was applied to construct a neighborhood graph, followed by Leiden clustering at resolutions of 0.4, 0.6, 1.0, and 1.4. For further analysis, Leiden clusters at a resolution of 0.4 were used for CD45^+^ and CD45^-^ cells, while a resolution of 1.0 was chosen for the pericyte-enriched fraction. Clusters containing fewer than 100 cells or those containing predominantly cells marked as doublets were removed. Additionally, one Fibroblast cluster was negative for Malat1 indicating a low quality of cells as previously shown^54^, and was therefore removed, as well. For the two-dimensional visualization UMAP was used. Cell-type annotation was performed by calculating the average expression of individual genes of all clusters and manually comparing them to known marker genes based on relevant publications^55,56^. Data was also automatically annotated to the Mouse Cell Atlas (MCA) using the scMCA R package^57^ (https://github.com/ggjlab/scMCA). Pseudotime analysis was performed by computing the diffusion map embedding, followed by the construction of the diffusion map based neighborhood graph (n_neighbors = 10). Differentially expressed genes (DEG) were calculated with a pseudo-bulk approach. To achieve this, UMI counts were aggregated over all cells for each individual cluster and for all individual mice. Resulting count tables were used in a subsequent DESeq2 analysis^58^. For the GSEA genes were ranked for their Wald-statistic which resulted from the DESeq2 analysis. This was conducted using the clusterProfiler^59^ and fgsea packages in R, utilizing MSigDBs^60,61^ hallmark gene sets^62^.

A total of 10 distinct cell clusters were manually identified within the CD45^-^ population, including ATIs, ATIIs, ECs, aCAP cells, secretory cells, ciliated cells, adventitial fibroblasts, fibroblasts, pericyte/SMC cells, and mesothelium (Figure 4b, Extended Data Fig. 4b). In the CD45^+^ population, 16 distinct cell clusters were identified, comprising B-cells, plasma cells, monocytes, transitional monocytes/macrophages, Cd300e^+^ macrophages, alveolar macrophages, interstitial macrophages, Cd101^+^ neutrophils, Ngp^+^ neutrophils, Treml4^+^ neutrophils, dendritic cells, T-cells, proliferating T-cells, NK cells, mast cells/basophils, and B cell like (Extended Data Fig. 4a, c). Neutrophil clusters were classified based on distinct marker expression: Cd101^+^ neutrophils were identified by *Cd101*, *Csf3r and Il1b*; Ngp^+^ neutrophils by *Ngp, Ltf*, and *Camp*; and Treml4^+^ neutrophils by *Treml4, Cx3cr1*, and *Clec4a1* (Extended Data Fig. 6a-c)^63^. Diffusion pseudotime was calculated for Neutrophils with “Cd101*+* neutrophils” and for Monocytes with “Monocyte” as root cells, to order the cells along the inferred trajectory. Unless otherwise indicated, standard parameters were used for all three calculations. Pseudotime analysis suggests that Treml4*+*neutrophils primarily originate from Cd101^+^ neutrophils and Ngp^+^ neutrophils (Extended Data Fig. 6b). Monocyte/macrophage clusters expressing the classical monocyte/macrophage marker *Cd68* were classified based on distinct marker expression: alveolar macrophages by *Plet1, Lpl, Hebp1*^64^; interstitial macrophages by *C1qa, C1qb, C1qc* ^64,65^; *Cd300e*+ macrophages by *Adgre4, Cd300e, Dusp16*; monocytes by *Fn1, F13a1, Ccr2*; and transitional monocyte/macrophage clusters, which co-express some markers of both monocytes and Cd300e+ macrophages, by *Nr4a1, Apoc2, Ace* (Extended Data Fig. 6d-f). Pseudotime analysis suggests a directional shift of monocytes, indicating a potential transition toward Cd300e^+^ macrophage states (Extended Data Fig. 6e). In the B cell–like cluster, most cells expressed the B cell marker *Cd19*; however, some cells within this cluster also expressed the monocyte marker *F13a1* and the T cell marker *Cd3e*.

Within pericytes/SMCs positive of CD45^-^ cluster of 723 cells, 349 cells originated from WT mice (both RA and CS groups combined), and only 66 cells exhibited *Cox4i2* expression.

MACS sorted Pericytes were pre-processed identically until and including the alignment step with STAR solo. Data was then imported to Scanpy where ambient RNA removal was performed with scAR Version 0.7.0. An adjusted protocol was used to increase the retained number of cells while making certain only high-quality cells were kept. Therefore, cell filtering was performed so that each cell incorporated between 1200 and 40000 UMIs, comprised less than 20 % mitochondrial reads and information of at least 600 expressed genes was present. Gene removal threshold was set to at least 20 cells expressing the gene. A subcluster containing no Malat1 expression at all, as well as the genes Gm42418 and AY036118 were removed altogether before further processing, since these genes represent rRNA contamination^66^ and influenced the embedding calculations heavily. Further computational processing of the data was equal to the whole lung cell samples. Differentially expressed genes for the GSEA were calculated with Scanpy’s rank_gene_groups function. Clusters were merged and annotated for their expression of *Cox4i2* and/or *Acta2*, marking the clusters as “Cox4i2^high^/Acta2^high^”, “Cox4i2^high^/Acta2^low^” or “Cox4i2^low^/Acta2^high^”. The scRNA-seq data have been deposited in NCBI’s Gene Expression Omnibus (GEO) and are accessible under GEO Series accession numbers GSE314716 and GSE314717.

### Cigarette smoke extract (CSE) preparation

100% CSE was prepared by bubbling cigarette smoke from one 3R4F cigarette (Kentucky Tobacco Research & Development Center) into 10 mL of RPMI (FCS-free) using a vacuum pump within one minute. After pH adjustment (7.4) using 2M NaOH, the medium was sterile filtered (0.22 µm) and diluted to the required concentration^48,67^. CSE was used immediately within 20 minutes for cell treatment.

### Isolation of pericyte enriched fraction

Cells were MACS sorted by labelling with biotin-conjugated antibodies against CD45 (#130-110-657, Miltenyi Biotec), CD31 (#130-111-353, Miltenyi Biotec), CD326 (#130-117-751, Miltenyi Biotec), and CD140a (#130-101-502, Miltenyi Biotec) for 15 minutes at 4°C. After labelling, cells were subjected to negative selection by incubating with anti-biotin microbeads for 15 minutes at 4°C. Following this step, the unlabelled cells were incubated with anti-mouse CD140b PE-Cy7 (#25-1402-82, Invitrogen) for 20 minutes at 4°C in the dark.

### Isolation of mouse primary pulmonary pericytes

Mouse lungs were digested at 37°C with dispase (2.5 U/mL, #4942078001, Merck), Liberase (100 µg/mL, #5401127001, Sigma-Aldrich), and DNase I (0.1 mg/mL, # A3778, A. Hartenstein GmbH) in MEM + 25 mM HEPES. The suspension was centrifuged (300 ×g, 5 min, 4°C), and RBCs were lysed with RBC Lysis Buffer (#11814389001, Sigma-Aldrich). After stopping lysis with 2% FBS, cells were counted and MACS-sorted. Cells were labeled with biotin-conjugated anti-CD31 (#130-111-353, Miltenyi Biotec, CD45 (#130-110-657, Miltenyi Biotec), CD326 (#130-117-751, Miltenyi Biotec), and CD140a (#130-101-502, Miltenyi Biotec), followed by depletion with anti-biotin microbeads. Unlabelled cells were then positively selected for CD140b (#130-128-936, Miltenyi Biotec) using biotin-conjugated antibodies and anti-biotin microbeads. CD140b^+^ pericytes were cultured in low-glucose DMEM with 2% FBS, Pericyte Supplements (ScienCell), and 1% Pen-Strep.

### Isolation of primary mouse pulmonary endothelial cells

Single-cell suspensions from mouse lungs were prepared as for pericytes. Cells were MACS sorted by labelling with biotin-conjugated anti-CD45, CD326, and CD140a (15 min, 4°C), then depleted using anti-biotin microbeads. Unlabelled cells were positively selected with anti-biotin CD31 microbeads.

### Isolation of primary mouse precapillary PASMCs

ACTA2^+^ PCs were isolated as previously described^14,68^. Briefly, cells were isolated from precapillary pulmonary arterial vessels that were filled with a mixture of 0.5% low-melting-point agarose and 0.5% Fe□O□ particles (Sigma-Aldrich). This approach allows selective collection of precapillary vessels, as Fe□O□ particles cannot pass through capillaries. Using this method, we previously confirmed that the isolated cells are ACTA2-positive^68^.

### Isolation of neutrophils from mouse bone marrow

Tibiae and femora from C57BL/6J mice were aseptically removed after death, and bone shafts were flushed with PBS (Ca_2_/Mg_2_-free) using a 24G needle. Collected bone marrow was centrifuged (300 ×g, 5 min, 4°C), and cells were counted with a LUNA FX7 cell counter. Neutrophils were isolated using the Miltenyi mouse neutrophil isolation kit.

### *In vitro* tube formation assay

Each μ-Slide Angiogenesis well (Ibidi) was coated with 10□μL of growth factor-reduced Matrigel (Corning) on ice and polymerized at 37□°C for 1 hour. Primary endothelial cells (3□×□10□/mL) in MV-2 basal medium (Promocell) with 0.4% heparin, 0.1% ascorbic acid, and 0.1% hydrocortisone were seeded at 15,000 cells/well. For co-culture, 3,000 pericytes were added. The plate was incubated at 37□°C, 5% CO_2_, and imaged hourly for 16–18 hours (EVOS M700, Thermo Fisher Scientific). Images at 7 hours were analysed with Image Pro.

### Measurement of mtROS with Mitosox

MtROS levels were evaluated in primary PASMCs, pericytes, and CMT cells after loading with 5□µM MitoSOX Red (ThermoFisher) for 20 minutes at 37°C. MitoSOX signal in cells was analysed using a BD Canto flow cytometer^69^ or STELLARIS Confocal Microscope (Leica Microsystems).

### Chemotaxis migration assay of neutrophils via indirect co-culturing of mouse pericytes

Isolated mouse pericytes were treated with 50 ng/mL TLR4 agonist (tlrl-3pelps, InvivoGen) or left untreated for 16 hours. Then, 300 µl of supernatant was transferred to a Costar 24-well plate’s bottom chamber, and 200,000 neutrophils were added to the transwell insert. After 16-18 hours at 37°C, neutrophil migration into the bottom chamber was assessed using the Cell Migration/Chemotaxis Assay Kit (Abcam).

### Cell death assay

Cell death was assessed using the Kinetic Apoptosis Kit (Abcam) following the manufacturer’s instructions. Images were captured using the EVOS M700 imaging system at 37°C and 5% CO□.

### Western blot

WB analysis was performed using a standard protocol^14^. The loading control β□actin was detected with anti□β□actin (Abcam; 1:10□000; #ab8226). Following primary antibodies were used: anti-COX4I2 (1:1000; #11463-1-AP; Proteintech), anti-ANG2 (dilution□1:250; #□PA5-27297; ThermoFisher Scientific), anti-VEGFR22 (dilution□1:1000; #□26415-1-AP; Proteintech) anti-β-actin (1:10000; #ab8226;). Western blotting with anti□COX4I2 antibodies revealed two distinct bands at ∼17–18□kDa in lung homogenates from both mouse and human samples, whereas cell lysates displayed a single band at the same molecular weight. Therefore, corresponding lung homogenates or cell lysates from *Cox4i2^-/-^*mice were used.

### Immunohistochemistry staining of nitrotyrosine residues

For counting nitrotyrosine, slides were incubated with anti-nitrotyrosine (1:100, #AB5411, MilliporeSigma). Detection utilized the ZytoChem Plus phosphatase polymer kit (Zytomed Systems GmbH,) with Warp Red Chromogen (Biocare Medical). CAT Hematoxylin (Biocare Medical) was used for counterstaining. Stained areas were quantified using Qwin software (Leica microsystem), standardized to the total tissue surface per image. As a negative control, immunohistochemistry staining for nitrotyrosine residues was performed after chemically reducing nitrotyrosines to aminophenols using sodium dithionite.

### Immunofluorescence staining

For CD45^+^ and CD3^+^ cell counting, sections were incubated overnight at 4°C with a primary antibody mix: anti-CD45 (1:500, #ab10559, Abcam) and anti-CD3 (1:150, #RBK024-05, Zytomed). The next day, sections were incubated for 1 hour at room temperature with Donkey Anti-Rabbit A555 (1:400, #32732, BioLegend) and Donkey Anti-Goat A647 (1:400, #405322, BioLegend). Cells were quantified using ImageJ’s manual cell counting plugin.

For dtTomato visualization, PCLS slides were incubated overnight at 4°C with anti-CD31 (1:200, #AF3628-SP, R&D Systems) and anti-alpha smooth muscle actin (ACTA2; 1:200, #A5228, Sigma-Aldrich) Co-detection of mRNA *COX4I2* and protein markers was detected using RNAScope Multiplex FLv2 kit (Bio□Techne). Human *COX4I2* mRNA (#570351-C3) was detected using the Opal Polaris 7-color manual IHC kit (Akoya Biosciences). Samples were further sequentially stained against PDGFRB (AB69506, Abcam; RRID:AB_1269704) and CSPG4 (HPA002951, Sigma; RRID:AB_1854449) using the Opal Polaris kit followed by overnight incubation with primary labelled ACTA2 (F3777, Sigma; RRID:AB_476977), and vWF-CF405M (A0082, Dako; RRID:AB_2315602) antibodies. Mix-n-Stain CF405M kit (Biotium) was used to label vWF antibody. NucSpot 750/780 (Invitrogen) was used as a nuclear counterstain and slides mounted with Prolong Diamond Antifade Mountant (Invitrogen). Negative controls were performed in parallel by omission of the hybridization probe and antibodies.

### Multiplex assay

A custom-made mouse magnetic bead-based multiplex assay was used to analyse the levels of selected inflammatory mediators in BALF, cell culture supernatant and human plasma according to the manufacturer’s protocol (R&D Systems). The assay was conducted using the Bio-Plex 200 instrument (Bio-Rad Laboratories GmbH) and analysed with the Bio-Plex Manager software (Bio-Rad Laboratories GmbH)^48^.

### Statistics

All *in vivo* experiments were performed using randomized allocation. Whenever feasible, investigators were blinded to group assignment during outcome assessments, including lung function measurements, haemodynamic evaluations, echocardiography, histological analyses, and FMT–µCT imaging. Statistical analyses were conducted using Student’s two-tailed *t*-test, one-way ANOVA, or two-way ANOVA, as appropriate. Data normality was assessed using a quantile–quantile (Q–Q) plot. Cell counts and results from Western blot, ELISA, and multiplex assays were log-transformed prior to analysis. Data derived from log-transformed variables are presented on a log10-scaled *y*-axis. All *p* values are reported in the figures; *p* ≥ 0.05 is considered not significant. Statistical analyses were performed using Prism 10 (GraphPad Software Inc.).

